# The Proteostasis Network is a Therapeutic Target in Acute Myeloid Leukemia

**DOI:** 10.1101/2024.09.24.614781

**Authors:** Kentson Lam, Yoon Joon Kim, Carlo M. Ong, Andrea Z. Liu, Fanny J. Zhou, Mary Jean Sunshine, Bernadette A. Chua, Silvia Vicenzi, Pierce W. Ford, Jie-Hua Zhou, Yuning Hong, Eric J. Bennett, Leslie A. Crews, Edward D. Ball, Robert A.J. Signer

**Affiliations:** Division of Regenerative Medicine, Department of Medicine, Stem Cell Discovery Center, Sanford Stem Cell Institute, Moores Cancer Center, University of California San Diego, La Jolla, CA 92093 USA; Division of Blood and Marrow Transplant, Department of Medicine, Moores Cancer Center, University of California San Diego, La Jolla, CA 92093 USA; School of Biological Sciences, Department of Cell and Developmental Biology, University of California at San Diego, La Jolla, CA 92093 USA; Department of Biochemistry and Chemistry, La Trobe Institute for Molecular Science, La Trobe University, Melbourne, VIC 3086 Australia

## Abstract

Oncogenic growth places great strain and dependence on the proteostasis network. This has made proteostasis pathways attractive therapeutic targets in cancer, but efforts to drug these pathways have yielded disappointing clinical outcomes. One exception is proteasome inhibitors, which are approved for frontline treatment of multiple myeloma. However, proteasome inhibitors are largely ineffective for treatment of other cancers, including acute myeloid leukemia (AML), although reasons for these differences are unknown. Here, we determined that proteasome inhibitors are ineffective in AML due to inability to disrupt proteostasis. In response to proteasome inhibition, AML cells activated HSF1 and autophagy, two key stem cell proteostasis pathways, to prevent unfolded protein accumulation. Inactivation of *HSF1* sensitized human AML cells to proteasome inhibition, marked by unfolded protein accumulation, activation of the PERK-mediated integrated stress response, severe reductions in protein synthesis, proliferation and cell survival, and significant slowing of disease progression and extension of survival *in vivo*. Similarly, combined autophagy and proteasome inhibition suppressed proliferation, synergistically killed AML cells, and significantly reduced AML burden and extended survival *in vivo*. Furthermore, autophagy and proteasome inhibition preferentially suppressed protein synthesis and induced apoptosis in primary patient AML cells, including AML stem/progenitor cells, without severely affecting normal hematopoietic stem/progenitor cells. Combined autophagy and proteasome inhibition also activated the integrated stress response, but surprisingly this occurred in a PKR-dependent manner. These studies unravel how proteostasis pathways are co-opted to promote AML growth, progression and drug resistance, and reveal that disabling the proteostasis network is a promising strategy to therapeutically target AML.

## INTRODUCTION

Proteostasis is regulated by pathways that control protein synthesis, folding, trafficking and degradation, and this network is frequently stressed in cancer^1,2^. Many oncogenic signaling pathways promote protein synthesis^3,4^, and malignant growth can exert significant demands on protein biosynthetic, folding, and turnover machinery^5,6^. Malignant genetic, environmental and metabolic stressors can also induce protein misfolding and damage that strain the ability of cancer cells to maintain proteostasis^7–9^, which can impair cell survival and proliferation^10–14^. To cope with proteostasis stress, some cancers activate and depend on stress response pathways, such as the heat shock response^10^ and the unfolded protein response (UPR)^7^. This dependency raised hopes that disabling stress response pathways or exacerbating stressed systems could overwhelm the resiliency of cancer cells to preserve proteostasis and drive their demise. Unfortunately, efforts to drug proteostasis pathways in cancer have yielded largely disappointing clinical outcomes^13,15,16^.

Despite these failures, proteasome inhibitors have proven highly effective for treating patients with multiple myeloma^13,17^. The proteasome is a key proteostasis pathway that degrades normal, damaged and misfolded proteins^18,19^. Bortezomib^20,21^ and carfilzomib^22–24^ are clinically approved proteasome inhibitors for frontline treatment of myeloma whose efficacy is partly linked to proteostasis disruption^20,21^. Despite success in myeloma, proteasome inhibitors have had largely disappointing results in patients with other cancers, including acute myeloid leukemia (AML)^25^. As a single agent, bortezomib has limited efficacy in relapse or refractory AML, although when used in combination with standard therapeutic regimens it modestly increased remission rates in older adults^15,25^. Another clinical trial did not show a benefit of adding bortezomib to standard care regimens in pediatric AML^15^. Why AML and other cancers tolerate proteasome inhibition is largely unknown.

Dichotomous outcomes in patients with myeloma and AML treated with proteasome inhibitors suggest there are important cancer-specific differences in proteostasis regulation. Recent advances indicate that myeloma is sensitive to proteasome inhibition due to underlying proteostasis stress in malignant plasma cells. Secretory plasma cells experience baseline ER stress associated with activation of the UPR, which shares eIF2α signaling with the integrated stress response (ISR)^26,27^. Phosphorylation of eIF2α^28^ suppresses protein synthesis^29^ and promotes preferential translation of ATF4^30^ that enhances expression of proteostasis factors^30,31^. However, if these adaptive changes are insufficient to resolve underlying stress, transcriptional output of the ISR can shift to drive apoptosis by promoting expression of CHOP^28,32^. Myeloma cells exhibit baseline ER stress and depend on adaptive ISR signaling at steady state; proteasome inhibition exacerbates ER stress and drives activation of a terminal ISR that kills myeloma cells^27^.

AML arises from transformation of hematopoietic stem (HSCs) and progenitor cells^33–35^ and depends on co-opted stem cell pathways^35–37^. The proteostasis network is uniquely configured in HSCs to restrict biogenesis and accumulation of misfolded and unfolded proteins^38–41^. HSCs exhibit low protein synthesis^38^, have minimal proteasome activity^39^, depend on autophagy^39^, and activate the heat shock response^40^ to preserve proteostasis and self-renewal. Mounting evidence has implicated autophagy in AML pathogenesis and metabolism^42,43^, but its impact on proteostasis is largely unknown. Heat shock factor 1 (HSF1), the master regulator of the heat shock response^44^, can promote AML growth^45,46^, and a subset of AML patients with low levels of phosphorylated-HSF1 had improved survival when bortezomib was added to standard of care treatments^47^, but whether HSF1 regulates proteostasis in AML is unknown. This raises the possibility that AML cells hijack stem cell mechanisms to preserve proteostasis and resist proteasome inhibition. The goal of this study was to determine how the proteostasis network is configured in AML, to determine mechanisms that confer resistance to proteasome inhibitors, and to establish if disrupting proteostasis can impair AML growth and progression.

## RESULTS

### Proteasome Inhibition Does Not Disrupt Proteostasis in AML

Efficacy of proteasome inhibitors in treatment of myeloma is linked to terminal UPR activation^27^. This suggests that proteostasis disruption is a prerequisite underlying sensitivity to proteasome inhibition. To test this, we treated three human myeloma cell lines with high (NCI-H929), moderate (U266), and low (RPMI 8226) proteasome inhibitor sensitivity^48,49^ with bortezomib for 24h and assessed impact on proteostasis by quantifying unfolded protein based on tetraphenylethene maleimide (TMI) fluorescence^50,39,41,51,52^. NCI-H929 and U266 cells exhibited significantly elevated TMI fluorescence in response to low- and high-dose bortezomib (Fig. 1A,B). In contrast, RPMI 8226 cells, which are less sensitive to bortezomib^48,49^, did not accumulate unfolded protein in response to high-dose bortezomib (Fig. 1C). These data link proteasome inhibitor sensitivity with unfolded protein accumulation in myeloma cells.

**Figure 1.**
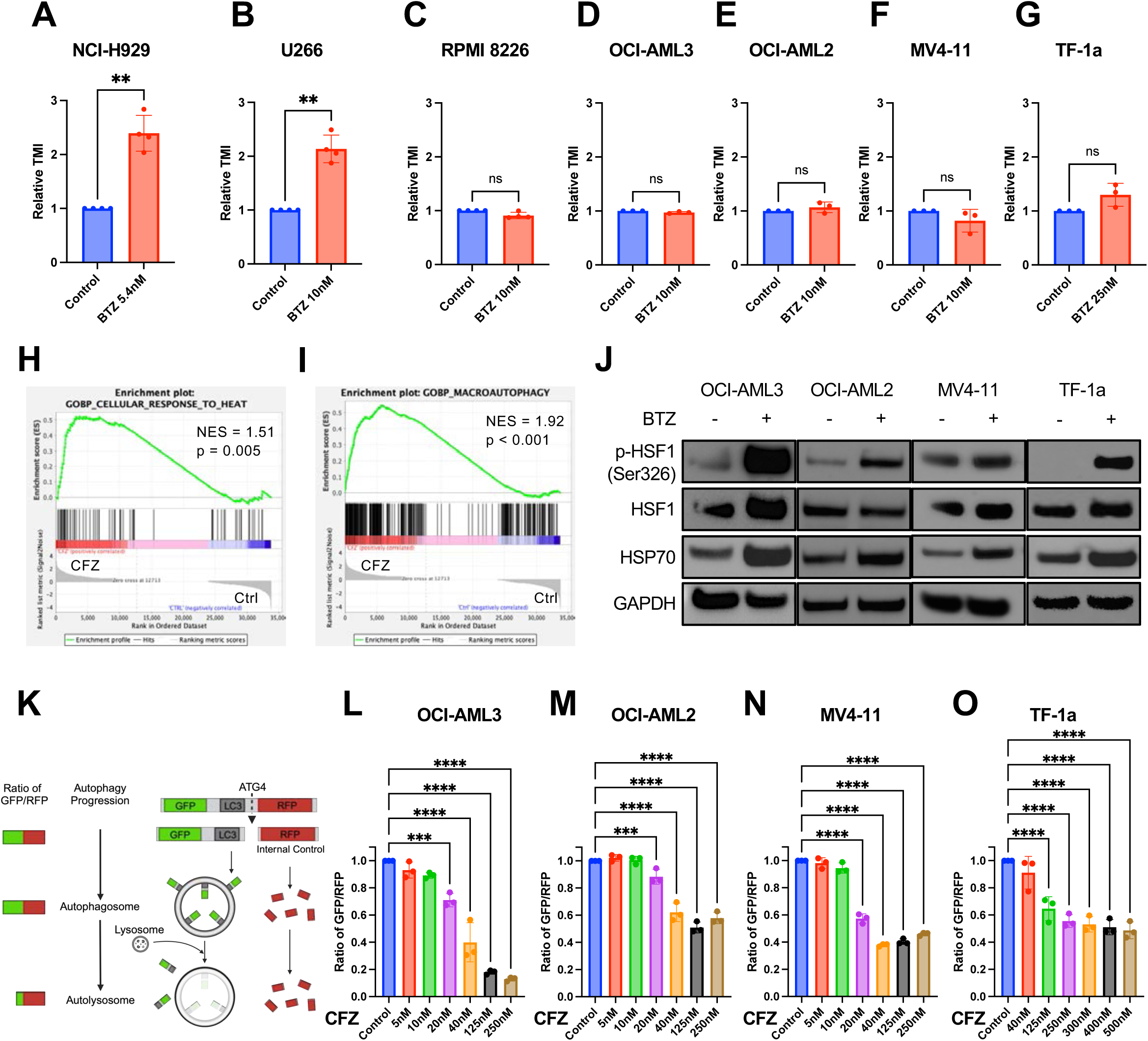
AML cells activate adaptive proteostasis pathways in response to proteasome inhibition. (A-C) Relative TMI fluorescence (unfolded protein content) detected by flow cytometry in (A) NCI-H929, (B) U266, and (C) RPMI 8226 human multiple myeloma cell lines treated with vehicle (PBS; control) or bortezomib (BTZ) at indicated concentrations for 24h (n=4). (D-G) Relative TMI fluorescence (unfolded protein content) detected by flow cytometry in (D) OCI-AML3, (E) OCI-AML2, (F) MV4-11, and (G) TF-1a human AML cell lines treated with vehicle (DMSO) or bortezomib at indicated concentrations for 24h (n=3). (H-I) Gene set enrichment analysis showing the (H) “cellular response to heat” and (I) “macroautophagy” gene sets in TF-1a cells treated for 24h with 125nM carfilzomib (CFZ) as compared to vehicle-treated controls (Ctrl; n=3). Normalized enrichment scores (NES) are shown. (J) Western blots for the indicated proteins involved in the heat shock response from OCI-AML3, OCI-AML2, MV4-11, and TF-1a human AML cell lines treated for 24h with vehicle (PBS; -) or their respective IC_50_ concentrations of bortezomib (BTZ; +). One of 3 representative blots shown. (K) Schematic of GFP-LC3-RFP reporter adapted from Kaizuka et al^59^, showing decrease in GFP to RFP ratio as autophagic flux increases. (L-O) Ratio of GFP:RFP fluorescence (inversely correlated to autophagic flux) measured by flow cytometry in (L) OCI-AML3, (M) OCI-AML2, (N) MV4-11, and (O) TF-1a human AML cell lines transduced with the RFP-GFP-LC3 reporter and treated with carfilzomib (CFZ) at the indicated concentrations (n=3). Data show individual replicates and mean ± standard deviation (SD) in (A-G, L-O). Statistical significance was assessed using a two-tailed paired student’s t-test (A-G) or ANOVA followed by Dunnett’s test relative to control (L-O); ∗p≤0.05, ∗∗p≤0.01, ∗∗∗p≤0.001, ∗∗∗∗p≤0.0001.

In contrast to myeloma, proteasome inhibitors are largely ineffective for treating AML^15,25^. We wondered if AML cells are refractory to proteasome inhibition because of failure to disrupt proteostasis. To test this, we treated four human AML cell lines (OCI-AML3, OCI-AML2, MV4-11, TF-1a) with high-dose bortezomib for 24h and quantified changes in unfolded protein. Bortezomib had no significant effect on unfolded protein content in any of the AML cell lines (Fig. 1D-G). This was not due to drug failure, as bortezomib suppressed proteasome activity in each cell line (Fig. S1A-D). These data indicate that proteasome inhibition is not sufficient to significantly disrupt proteostasis in AML cells.

### AML Cells Activate Adaptive Proteostasis Pathways in Response to Proteasome Inhibition

When one proteostasis pathway is disrupted, adaptive changes can rebalance the system to prevent proteostasis collapse^2,53^ ^18,54^. This raised the question of whether AML cells reconfigure their proteostasis network in response to proteasome inhibition to prevent accumulation of unfolded protein. To test this, we performed RNA-sequencing on TF-1a cells treated with high-dose carfilzomib. Proteasome inhibition resulted in substantial transcriptional changes in TF-1a cells (Fig. S1E-G). Gene set enrichment analysis suggested that at least two proteostasis pathways were activated in response to proteasome inhibition – the heat shock response and autophagy (Fig. 1H,I).

To complement RNA-sequencing data, we sought to determine if proteasome inhibition activated HSF1 in AML cells. Activated HSF1 promotes transcription of heat shock proteins and other factors that help restore proteostasis^55,56^. 24h-Bortezomib treatment substantially increased phosphorylated (Ser326) HSF1 – a marker of activation^57^ – across all AML cell lines (Fig. 1J). We also observed accumulation of HSP70, a key transcriptional target of HSF1^58^, in response to bortezomib (Fig. 1J). These data indicate that proteasome inhibition induces activation of HSF1 and the heat shock response in AML cells.

RNA-sequencing also suggested that AML cells activate autophagy in response to proteasome inhibition (Fig. 1I). We confirmed this by quantifying autophagic flux. We transduced all four AML cell lines with a dual-fluorescent autophagy reporter (LC3-GFP-RFP; Fig. 1K)^59^. Activation of autophagy induces cleavage of RFP by the autophagy-related protease ATG4 and relocalization of GFP-LC3 to autophagosomes^59^. Fusion of autophagosomes with lysosomes quenches and degrades GFP^59,60^, while cleaved RFP is stable in the cytoplasm and serves as an internal control. Declines in GFP:RFP indicate increased autophagic flux^59^. We treated AML cells with carfilzomib for 24h and found that GFP:RFP was reduced in a dose-dependent manner across all four AML cell lines (Fig. 1L-O). Bortezomib had similar effects (Fig. S1H-K) indicating that these effects are a conserved feature of proteasome inhibition. Thus, proteasome inhibition increases autophagic flux in AML cells.

### HSF1 Promotes AML Proteostasis, Viability and Proliferation in Response to Proteasome Inhibition

Proteasome inhibitors induced the heat shock response in AML cells (Fig. 1H,J), but whether this contributed to preventing proteostasis disruption and drug resistance was unknown. To test this, we used CRISPR/Cas9 to delete *HSF1*. Two gRNAs targeting different exons were used to generate distinct *HSF1*-deficient clones for each of the four AML cell lines. *HSF1* deletion was validated by sequencing (not shown) and confirmed by western blot at steady state and in response to bortezomib (Fig. 2A). *HSF1*-deficiency on its own did not typically affect unfolded protein abundance in AML cells (Fig. 2B-D, S2E-H). However, in contrast to wildtype controls, all four *HSF1*-deficient AML cell lines exhibited significant accumulation of unfolded protein following 24h bortezomib treatment (Fig. 2B-E, S2E-H). These data indicate that HSF1 helps preserve AML proteostasis by preventing accumulation of unfolded protein in response to proteasome inhibition.

**Figure 2.**
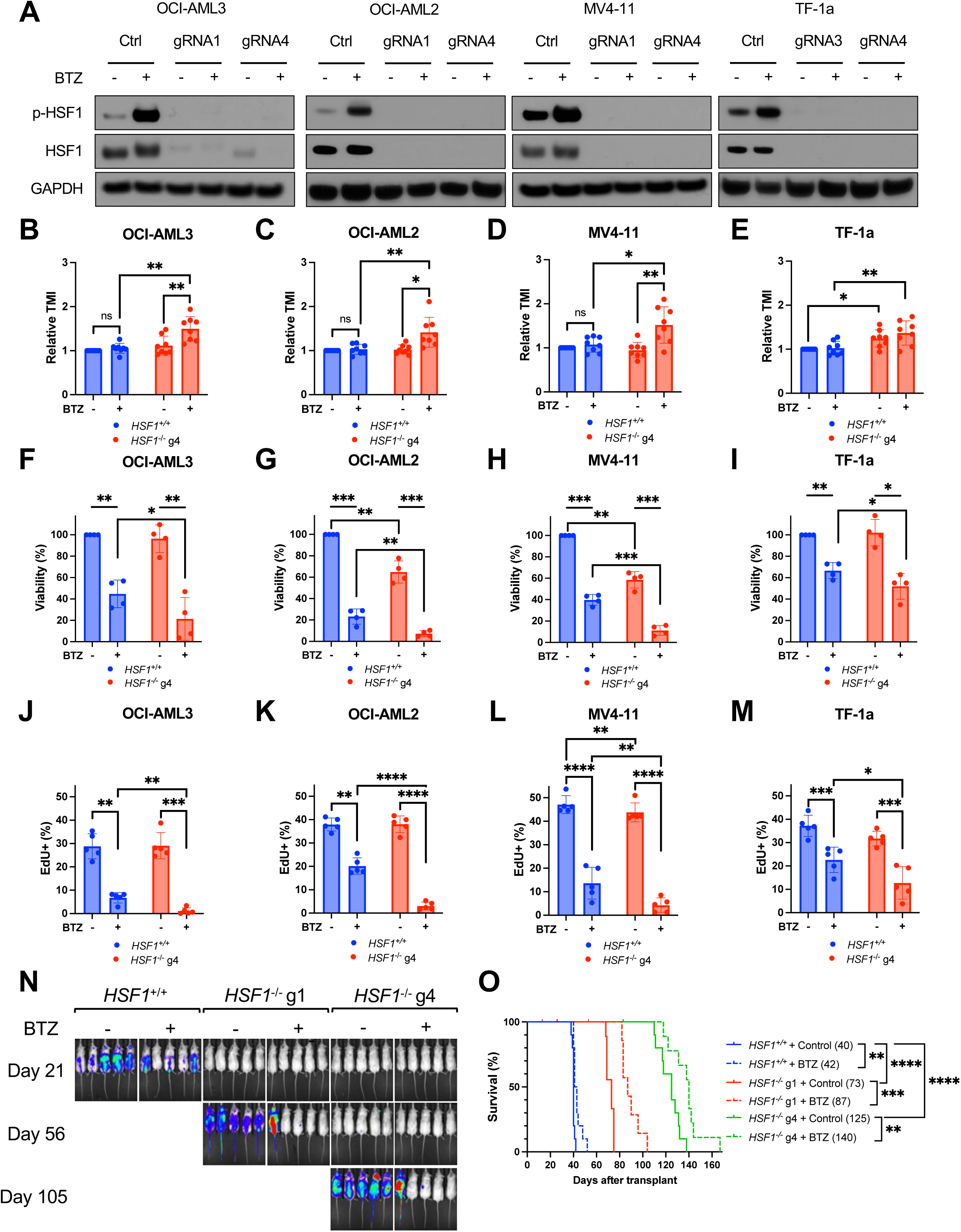
*HSF1* deletion sensitizes human AML cells to proteasome inhibition *in vitro* and *in vivo*. (A) Western blots assessing HSF1 expression and activation (Ser326 phosphorylation) in wildtype and *HSF1*-deleted OCI-AML3, OCI-AML2, MV4-11, and TF-1a human AML cell lines. Cells were treated for 24h with vehicle (PBS; -) or their respective IC_50_ concentrations of bortezomib (BTZ; +). Two distinct *HSF1*-deficient clones (generated with the indicated gRNAs) are shown for each cell line. (B-E) Relative TMI fluorescence (unfolded protein content) detected by flow cytometry in wildtype (*HSF1*^+/+^) and *HSF1*^-/-^ (B) OCI-AML3, (C) OCI-AML2, (D) MV4-11, and (E) TF-1a human AML cell lines treated for 24h with vehicle (PBS; -) or their respective IC_50_ concentrations of bortezomib (BTZ; +; n=8). (F-I) Frequency of viable cells (measured by MTT) in wildtype (*HSF1*^+/+^) and *HSF1*^-/-^ (F) OCI- AML3, (G) OCI-AML2, (H) MV4-11, and (I) TF-1a human AML cell lines treated for 24h with vehicle (PBS; -) or their respective IC_50_ concentrations of bortezomib (BTZ; +; n=4). (J-M) Frequency of dividing EdU^+^ (following 1h pulse) wildtype (*HSF1*^+/+^) and *HSF1*^-/-^ (J) OCI- AML3, (K) OCI-AML2, (L) MV4-11, and (M) TF-1a cells treated with vehicle (PBS; -) or their respective IC_50_ concentrations of bortezomib (BTZ; +; n=5). (N,O) (N) Representative bioluminescence images and (O) overall survival of NSG mice xenotransplanted with wildtype (*HSF1*^+/+^) or *HSF1*^-/-^ OCI-AML3 cells (transduced with luciferase). Mice were treated with PBS (-) or 0.5 mg/kg bortezomib (+). Treatment was initiated 5 days after transplant and drug was administered weekly for 2 consecutive days followed by 5 days with no treatment. (N) Images are at the indicated times after transplantation. (O) Median survival (days) is shown in parentheses (n=10/group). Data show individual replicates and mean ± SD in (B-M). Statistical significance was assessed using a two-tailed paired student’s t-test (B-M) or log-rank test (O); ∗p≤0.05, ∗∗p≤0.01, ∗∗∗p≤0.001, ∗∗∗∗p≤0.0001.

We further examined if HSF1 conferred AML resistance to proteasome inhibition by testing effects on viability and proliferation. Control and *HSF1*-deficient AML cell lines were cultured in the presence of bortezomib at their IC_50_ concentrations (Fig. S2A-D) or vehicle for 24h and viability was quantified via MTT assay. *HSF1*-deficiency reduced viability in OCI-AML2 and MV4-11 AML cells but had no significant effects on OCI-AML3 or TF-1a cells (Fig. 2F-I, S2I-L), suggesting a potential context specific role for HSF1 in AML cells at steady state. However, *HSF1*-deficiency significantly sensitized all four human AML cell lines to proteasome inhibition as viability was significantly lower in all four bortezomib-treated *HSF1*-deficient cell lines as compared to bortezomib-treated controls (Fig. 2F-I, S2I-L). Furthermore, anti-proliferative effects of proteasome inhibition were significantly enhanced by *HSF1*-deficiency. Although *HSF1*-deficiency had minimal effects on baseline AML cell proliferation, EdU incorporation was significantly lower in all four bortezomib-treated *HSF1*-deficient AML cell lines as compared to bortezomib-treated controls (Fig. 2J-M, S2M-P). We confirmed that effects of *HSF1*-deficiency were generally conserved between individual AML cell line clones, although in rare cases small differences were observed (Fig. S2).

These data indicate that HSF1 has minimal effects on AML proteostasis, viability and proliferation at steady state, but that it confers broad protection against proteasome inhibition by preventing accumulation of unfolded protein, enhancing survival and promoting proliferation irrespective of underlying genetic mutations across multiple human AML cell lines.

### HSF1 Promotes AML Progression and Confers Resistance to Proteasome Inhibitors *In Vivo*

To examine the impact of HSF1 on AML *in vivo*, we transduced wildtype or *HSF1*-deficient OCI-AML3 cells with a luciferase reporter^61^ and xenotransplanted 5×10^5^ cells into immunodeficient NSG mice^62^. Five days after transplant, half the mice were treated with bortezomib (0.5 mg/kg; 2 days on, 5 days off) or vehicle for the duration of the experiment. AML progression was monitored by *in vivo* bioluminescence imaging. Consistent with prior reports^63^, bortezomib had minimal effects on AML progression and overall survival, as leukemia burden was only modestly reduced in bortezomib-treated mice (Fig. 2N) and median survival was only extended from 40 to 42 days (Fig. 2O). *HSF1*-deficiency reduced AML burden and extended median survival from 40 to 73 (clone g1) or 125 (clone g4) days (Fig. 2N,O). Bortezomib treatment further reduced leukemia burden and extended median survival in *HSF1*-deficient AML xenograft recipients from 73 to 87 days and 125 to 140 days, respectively (Fig. 2N,O). *HSF1*-deficiency also extended survival and conferred enhanced sensitivity to bortezomib in more aggressive TF-1a (*TP53*- mutant) xenografts (Fig. S2Q-S). These data indicate that HSF1 promotes AML progression and limits the efficacy of proteasome inhibitors *in vivo*.

### Combined Proteasome and Autophagy Inhibition Synergistically Disrupts AML Cell Proteostasis, Viability and Proliferation

Although proteasome is the canonical protein degradation machinery^64,65^, autophagy can regulate protein turnover - especially in response to proteasome inhibition^18,66^. HSCs increase autophagy in response to proteasome inhibition, and disabling both proteasome and autophagy is required for increasing misfolded/unfolded proteins within HSCs^39^. AML cells also increase autophagy in response to proteasome inhibition (Fig. 1L-O, S1H-K), but whether this contributes to preserving proteostasis, viability and proliferation is unknown.

To test this, we cultured four AML cell lines with the proteasome inhibitor carfilzomib, the autophagy inhibitor Lys05^67^ (Fig. S3A), the combination of the two inhibitors, or vehicle for 24h and quantified unfolded protein, viability and proliferation. Drug concentrations were based on empirically determined IC_50_ concentrations (Fig. S3B-I). Carfilzomib or Lys05 treatment on their own did not significantly impact unfolded protein (Fig. 3A-D). However, combined carfilzomib/Lys05 treatment significantly increased unfolded protein by ∼65%-170% across all four AML cell lines (Fig. 3A-D). Combined carfilzomib/Lys05 treatment also reduced viability by >86% (Fig. 3E-H) and proliferation by >95% (Fig. 3I-L) in all four AML cell lines. Synergy analyses using the Chou-Talalay method^68^ and SynergyFinder+^69^ indicated that combined carfilzomib/Lys05 treatment exhibited synergistic effects on viability (Fig. S3J,K; Combination Indexes <1 and Synergy Scores >10). These data indicate that autophagy confers AML cells with resistance to proteasome inhibitors by preventing proteostasis disruption, and combined proteasome and autophagy inhibition severely impairs AML proteostasis, survival and growth.

**Figure 3.**
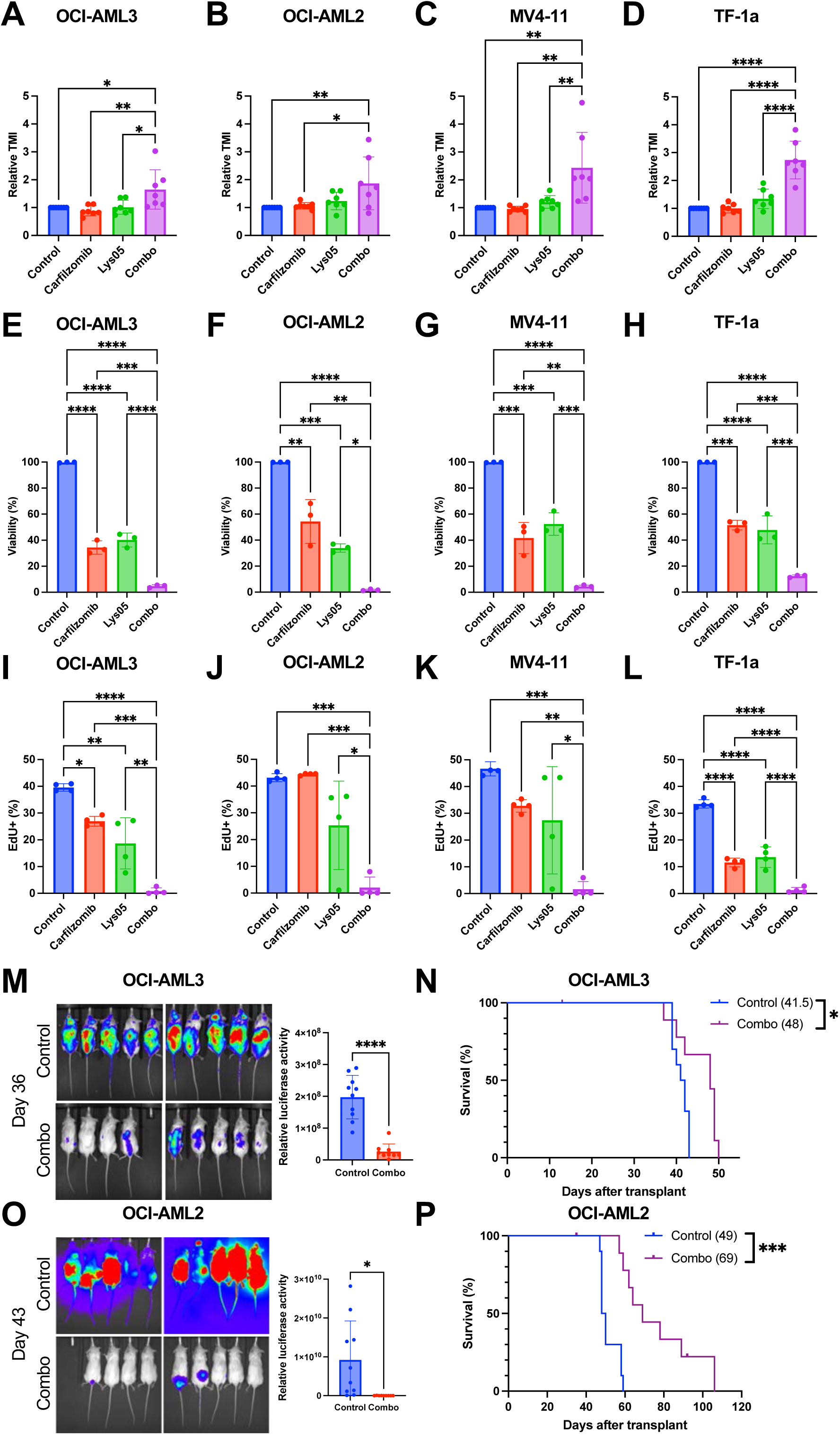
Combined proteasome and autophagy inhibition synergistically disrupts AML proteostasis, proliferation and survival *in vitro* and impairs AML progression *in vivo*. (A-D) Relative TMI fluorescence (unfolded protein content) detected by flow cytometry in (A) OCI-AML3, (B) OCI-AML2, (C) MV4-11, and (D) TF-1a human AML cell lines treated for 24h with DMSO (Control), carfilzomib, Lys05, or the combination of carfilzomib and Lys05 (Combo). Carfilzomib and Lys05 were used at their respective IC_50_ concentrations for each cell line (n=7). (E-H) Frequency of viable cells (measured by MTT) in (E) OCI-AML3, (F) OCI-AML2, (G) MV4-11, and (H) TF-1a human AML cell lines treated for 24h with DMSO (Control), carfilzomib, Lys05, or the combination of carfilzomib and Lys05 (Combo). Carfilzomib and Lys05 were used at their respective IC_50_ concentrations for each cell line (n=3). (I-L) Frequency of dividing EdU^+^ (following 1h pulse) (I) OCI-AML3, (J) OCI-AML2, (K) MV4-11, and (L) TF-1a cells treated for 24h with DMSO (Control), carfilzomib, Lys05, or the combination of carfilzomib and Lys05 (Combo). Carfilzomib and Lys05 were used at their respective IC_50_ concentrations for each cell line (n=4). (M-P) Representative bioluminescence images, bioluminescence quantification, and overall survival of NSG mice xenotransplanted with (M,N) OCI-AML3 cells (transduced with luciferase) or (O,P) OCI-AML2 cells (transduced with luciferase). Mice were treated with PBS (Control) or 0.5mg/kg bortezomib and 30mg/kg Lys05 (Combo). Treatment was initiated 7 days after transplant and drug was administered weekly for 2 consecutive days followed by 5 days with no treatment. (M,O) Images are at the indicated times after transplantation. (N,P) Median survival (days) is shown in parentheses (n=9-10/group). Data show individual replicates and mean ± SD in (A-M, O). Statistical significance was assessed using an ANOVA followed by Tukey’s test (A-L), a two-tailed unpaired t-test (M,O), or log-rank test (N,P); ∗p≤0.05, ∗∗p≤0.01, ∗∗∗p≤0.001, ∗∗∗∗p≤0.0001.

### Combined Proteasome and Autophagy Inhibition Impairs AML Progression and Extends Survival *In Vivo*

To test the efficacy of combined proteasome/autophagy inhibition *in vivo*, luciferase-transduced OCI-AML3 and OCI-AML2 cells were xenotransplanted into NSG mice. Seven days after transplant, mice were treated with 0.5 mg/kg bortezomib (intravenous) and 30 mg/kg Lys05 (intraperitoneal; 2 days on, 5 days off) or vehicle for the duration of the experiment. Combined bortezomib/Lys05 treatment reduced AML burden and significantly extended survival in both OCI-AML3 and OCI-AML2 xenografts (Fig. 3M-P). Combined bortezomib/Lys05 treatment extended median survival from 41.5 to 48 days (OCI-AML3; Fig. 3N) and 49 to 69 days (OCI- AML2; Fig. 3P). These data indicate that combined proteasome/autophagy inhibition impairs progression of AML xenografts and significantly extends survival.

### Proteostasis Disruption Activates a Terminal Integrated Stress Response in AML Cells

To test if proteostasis disruption contributed to impaired AML growth and survival and to gain additional mechanistic insights, we performed RNA-sequencing on carfilzomib-treated *HSF1*- deficient and wildtype TF-1a cells (Fig. S4A,B). We also performed RNA-sequencing on combined carfilzomib/Lys05- and vehicle-treated wildtype TF-1a cells (Fig. S4C,D). Strikingly, pathway analyses revealed that EIF2 signaling was the most upregulated pathway in *both* carfilzomib-treated *HSF1*-deficient and combined carfilzomib/Lys05-treated TF-1a cells as compared to their respective controls (Fig. 4A,B). EIF2 promotes translation initiation and is essential for protein synthesis^70^. EIF2 activity can be attenuated by the ISR through EIF2α phosphorylation^28^, which reduces protein synthesis^29^ and promotes preferential translation of ATF4^30^. Consistent with changes in EIF2 signaling, gene set enrichment analysis revealed significant upregulation of ISR signaling in both carfilzomib-treated *HSF1*-deficient and carfilzomib/Lys05-treated TF-1a cells compared to their respective controls (Fig. 4C,D).

**Figure 4.**
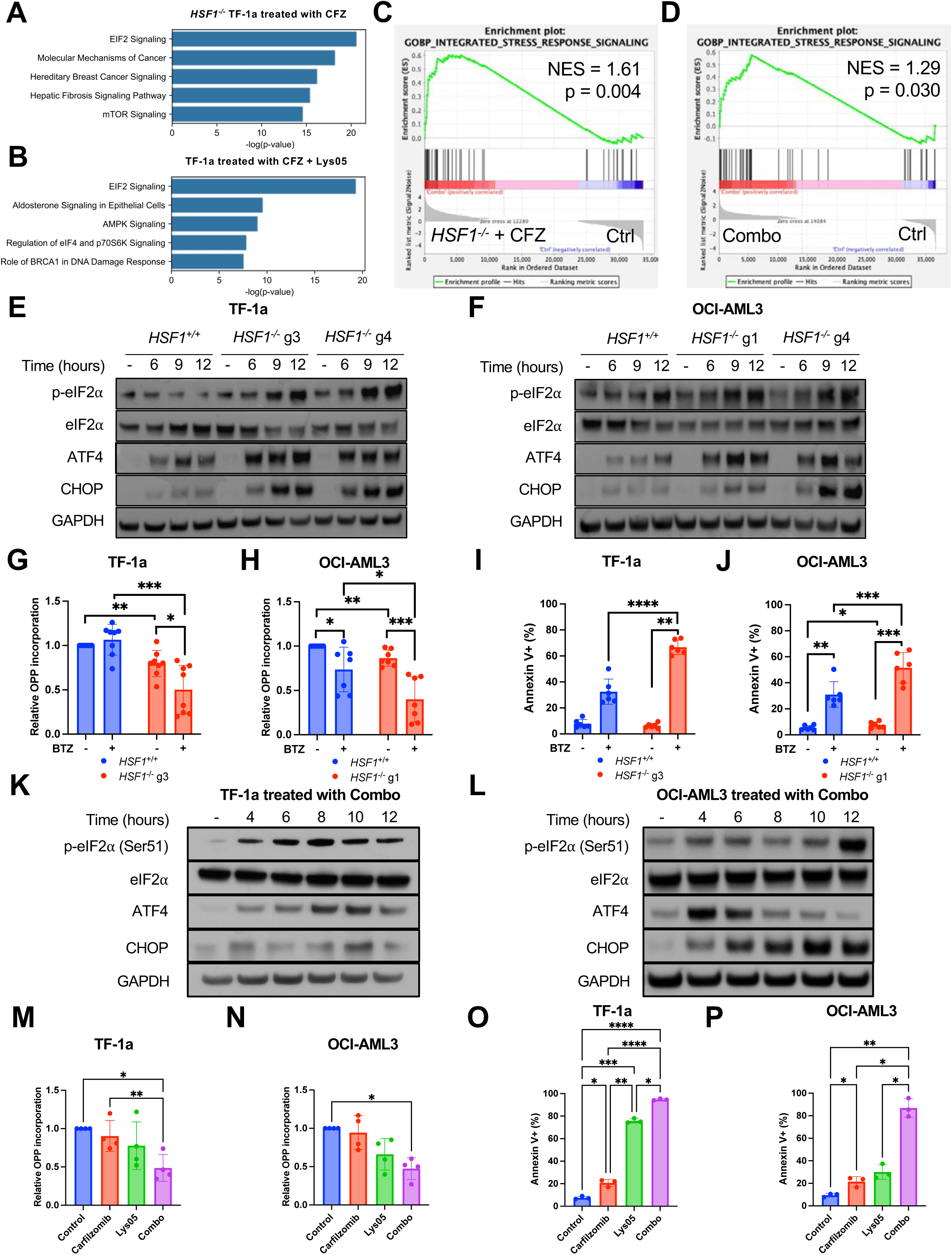
Proteostasis disruption activates a terminal ISR in human AML cells. (A-B) The top 5 upregulated canonical pathways (Qiagen Ingenuity Pathway Analysis) in (A) *HSF1*^-/-^ TF-1a cells treated for 24h with 125nM carfilzomib (CFZ) and (B) TF-1a cells treated for 24h with 125nM CFZ and 7.5µM Lys05 as compared to untreated wildtype TF-1a cells. (C-D) Gene set enrichment analysis showing upregulation of the “integrated stress response signaling” gene set in (C) *HSF1*^-/-^ TF-1a cells treated for 24h with 125nM carfilzomib (CFZ) and (D) TF-1a cells treated for 24h with 125nM CFZ and 7.5µM Lys05 (Combo) as compared to DMSO-treated (Ctrl) wildtype TF-1a cells. Normalized enrichment scores (NES) are shown. (E-F) Western blots showing time course expression of various proteins involved in the ISR in wildtype (*HSF1*^+/+^) and *HSF1*^-/-^ (E) TF-1a and (F) OCI-AML3 cells treated with their respective IC_50_ concentrations of bortezomib for the indicated times. (G-H) Relative O-propargyl-puromycin (OPP) incorporation (protein synthesis) in wildtype (*HSF1*^+/+^) and *HSF1*^-/-^ (G) TF-1a (n=8) and (H) OCI-AML3 (n=7) cells treated for 24h with vehicle (PBS; -) or their respective IC_50_ concentrations of bortezomib (BTZ; +). (I-J) Frequency of apoptotic Annexin V^+^ wildtype (*HSF1*^+/+^) and *HSF1*^-/-^ (I) TF-1a (n=6) and (J) OCI-AML3 (n=6) cells treated for 24h with vehicle (PBS; -) or their respective IC_50_ concentrations of bortezomib (BTZ; +). (K-L) Western blots showing time course expression of various proteins involved in the ISR in (K) TF-1a and (L) OCI-AML3 treated with the combination of carfilzomib and Lys05 (Combo) at their respective IC_50_ concentrations. (M-N) Relative OPP incorporation (protein synthesis) in (M) TF-1a and (N) OCI-AML3 treated for 24h with DMSO (Control), carfilzomib, Lys05, or the combination of carfilzomib and Lys05 (Combo). Carfilzomib and Lys05 were used at their respective IC_50_ concentrations for each cell line (n=4). (O-P) Frequency of apoptotic Annexin V^+^ (O) TF-1a and (P) OCI-AML3 treated for 24h with DMSO (Control), carfilzomib, Lys05, or the combination of carfilzomib and Lys05 (Combo). Carfilzomib and Lys05 were used at their respective IC_50_ concentrations for each cell line (n=3). Data show individual replicates and mean ± SD in (G-J, M-P). Statistical significance was assessed using a two-tailed paired student’s t-test (G-J) or ANOVA followed by Tukey’s test (M-P); ∗p≤0.05, ∗∗p≤0.01, ∗∗∗p≤0.001, ∗∗∗∗p≤0.0001.

To confirm ISR activation, we performed time-course western blots to assess EIF2α- phosphorylation, ATF4 and CHOP abundance^32^. Bortezomib treatment on its own did not induce substantial EIF2α-phosphorylation in either TF-1a or OCI-AML3 cells, although ATF4 and CHOP abundance modestly increased (Fig. 4E,F). In contrast, bortezomib treatment induced robust EIF2α-phosphorylation as well as ATF4 and CHOP expression in *HSF1*-deficient TF-1a and OCI-AML3 cells (Fig. 4E,F), confirming ISR activation.

We also quantified effects on protein synthesis (O-propargyl-puromycin (OPP) incorporation^71^) and apoptosis (Annexin V). Consistent with nominal ISR activation, bortezomib on its own had minimal effects on protein synthesis. In contrast, bortezomib significantly reduced OPP incorporation in all four *HSF1*-deficient AML cell lines compared to both vehicle-treated *HSF1*- deficient controls and bortezomib-treated wildtype AML cells (Fig. 4G,H, S4E,F). In addition, bortezomib induced apoptosis in ∼50-80% of all four *HSF1*-deficient AML cell lines, and frequency of Annexin V^+^ cells was significantly higher in bortezomib-treated *HSF1*-deficient AML cells compared to bortezomib-treated controls (Fig. 4I,J, S4G,H). These data indicate that HSF1 largely protects AML cells from ISR activation in response to proteasome inhibition, but in the absence of HSF1, proteasome inhibition leads to rapid, robust ISR activation in AML cells.

Combined bortezomib/Lys05 treatment also induced rapid EIF2α-phosphorylation and upregulation of ATF4 and CHOP (Fig. 4K,L). These molecular changes were accompanied by ∼50% reduction in protein synthesis (Fig. 4M,N; Fig. S4I,J) and >85% apoptosis (Fig. 4O,P, S4K,L) in all four AML cell lines. These data indicate that combined proteasome/autophagy inhibition leads to rapid, robust ISR activation in AML cells.

### PERK Promotes ISR Activation in *HSF1*-deficient AML Cells in Response to Proteasome Inhibition

ISR signaling can be induced by four known kinases: HRI (*EIF2AK1*), PKR (*EIF2AK2*), PERK (*EIF2AK3*) and GCN2 (*EIF2AK4*)^72–75^. To determine which kinase promotes ISR activation in response to proteostasis disruption in AML cells, we treated *HSF1*-deficient and wildtype TF-1a cells with carfilzomib or vehicle for 24h and performed RNA-sequencing (Fig. S5A). Of the four kinases, only *EIF2AK3* (PERK) was specifically upregulated in carfilzomib-treated *HSF1*- deficient AML cells (Fig. 5A). PERK is typically activated in response to ER stress, which can be induced by accumulation of unfolded protein within the ER^76^. Carfilzomib was sufficient to induce upregulation of the “response to endoplasmic reticulum stress” gene set in TF-1a cells (Fig. 5B), but it was further upregulated in carfilzomib-treated *HSF1*-deficient TF-1a cells (Fig. 5C). Furthermore, carfilzomib-treated *HSF1*-deficient TF-1a cells significantly upregulated the “PERK mediated unfolded protein response” gene set (Fig. 5D).

**Figure 5.**
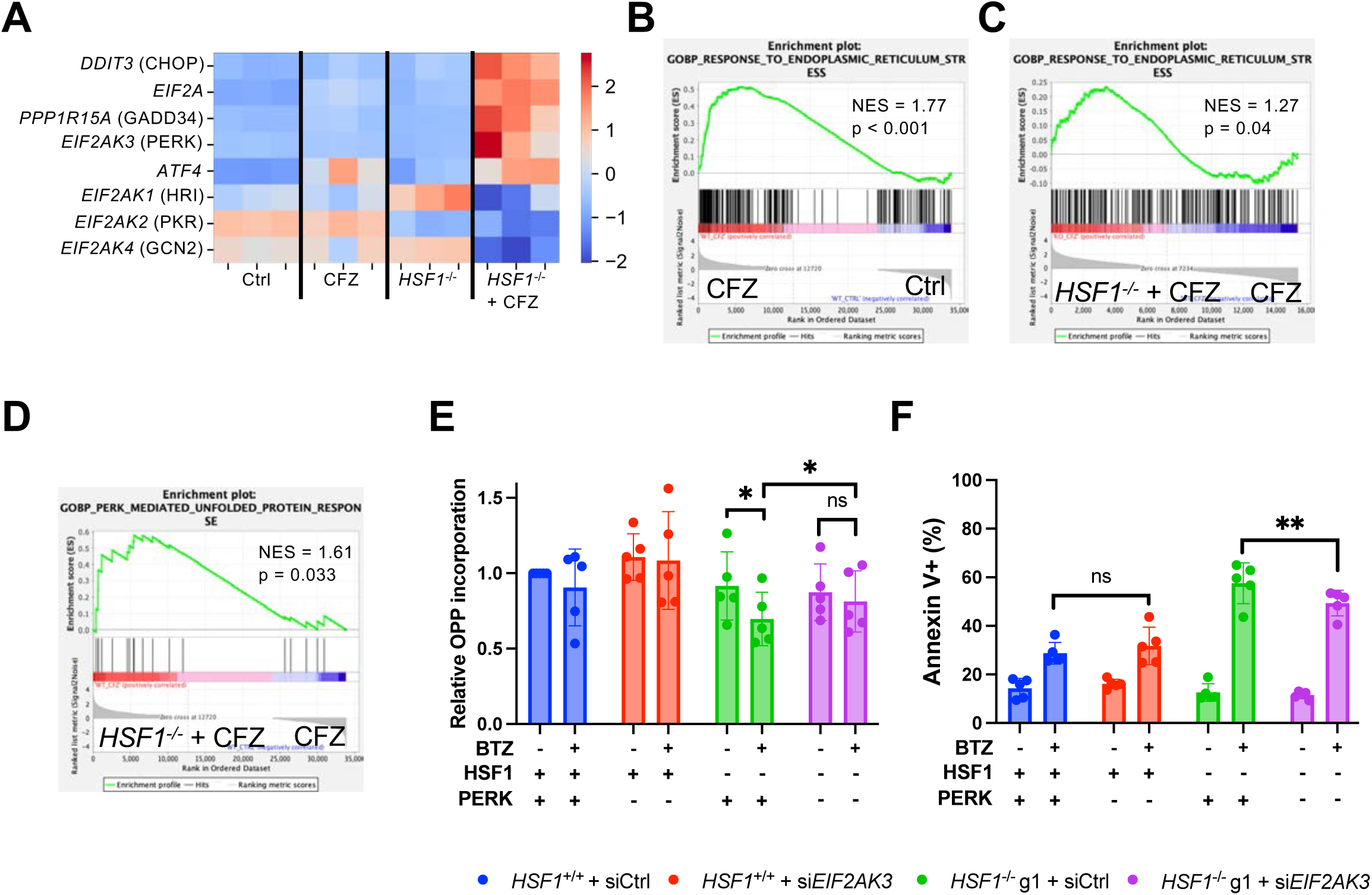
PERK promotes ISR activation in *HSF1*-deficient AML cells in response to proteasome inhibition. (A) Heatmap depicting the relative expression (based on RNA-sequencing; n=3) of some ISR genes in wildtype and *HSF1*^-/-^ TF-1a cells treated for 24h with vehicle (PBS; Ctrl) or 125nM carfilzomib (CFZ). (B-D) Gene set enrichment analysis showing enrichment of the (B,C) “endoplasmic reticulum stress” and (D) “PERK mediated unfolded protein response” gene sets in (B) TF-1a cells treated for 24h with 125nM carfilzomib (CFZ) as compared to vehicle treated controls (Ctrl) and (C,D) *HSF1*^-/-^ TF-1a cells treated for 24h with 125nM carfilzomib (*HSF1*^-/-^ + CFZ) as compared to similarly treated wildtype (*HSF1*^+/+^) TF-1a (CFZ). Normalized enrichment scores (NES) are shown. (E) Relative OPP incorporation (protein synthesis) and (F) frequency of apoptotic Annexin V^+^ cells in wildtype (*HSF1*^+/+^) and *HSF1*^-/-^ OCI-AML3 transfected with control (siCtrl) or PERK-targeted siRNA (si*EIF2AK3*) treated for 24h with vehicle (PBS) or 7.5nM bortezomib (BTZ) (n=5). Data show individual replicates and mean ± SD in (E,F). Statistical significance was assessed using a two-tailed paired student’s t-test (E,F). ∗p≤0.05, ∗∗p≤0.01.

To test if PERK had a role in driving effects of proteasome inhibition in *HSF1*-deficient AML cells, we used *EIF2AK3*-targeted siRNA to knock down PERK in wildtype and *HSF1*-deficient OCI-AML3 cells (Fig. S5B). PERK knockdown had no significant effects on protein synthesis or apoptosis in control or bortezomib-treated OCI-AML3 cells (Fig. 5E,F). However, PERK knockdown significantly attenuated the reduction in protein synthesis (Fig. 5E) and the induction of apoptosis (Fig. 5F) caused by bortezomib in *HSF1*-deficient AML cells. These data suggest that HSF1 protects AML cells from proteasome inhibition by preserving proteostasis and preventing UPR activation. In the absence of HSF1, proteasome inhibition causes AML cells to accumulate unfolded protein, activate the PERK-mediated UPR and trigger the ISR that suppresses protein synthesis, reduces proliferation and induces apoptosis to impair AML growth and progression.

### PKR Promotes ISR Activation in AML Cells in Response to Combined Proteasome and Autophagy Inhibition

Next, we sought to determine if ISR activation downstream of combined proteasome/autophagy inhibition was also driven by PERK. We deleted *EIF2AK3* from OCI-AML3 cells (Fig. S6A) and examined effects of combined carfilzomib/Lys05 treatment on wildtype and *EIF2AK3*^-/-^ AML cells. Carfilzomib/Lys05 treatment severely suppressed protein synthesis and induced apoptosis in OCI-AML3 cells – consistent with ISR activation. However, PERK-deficiency did not significantly rescue these phenotypes (Fig. 6A,B). Thus, PERK is not required to induce ISR effects following combined proteasome/autophagy inhibition in AML.

**Figure 6.**
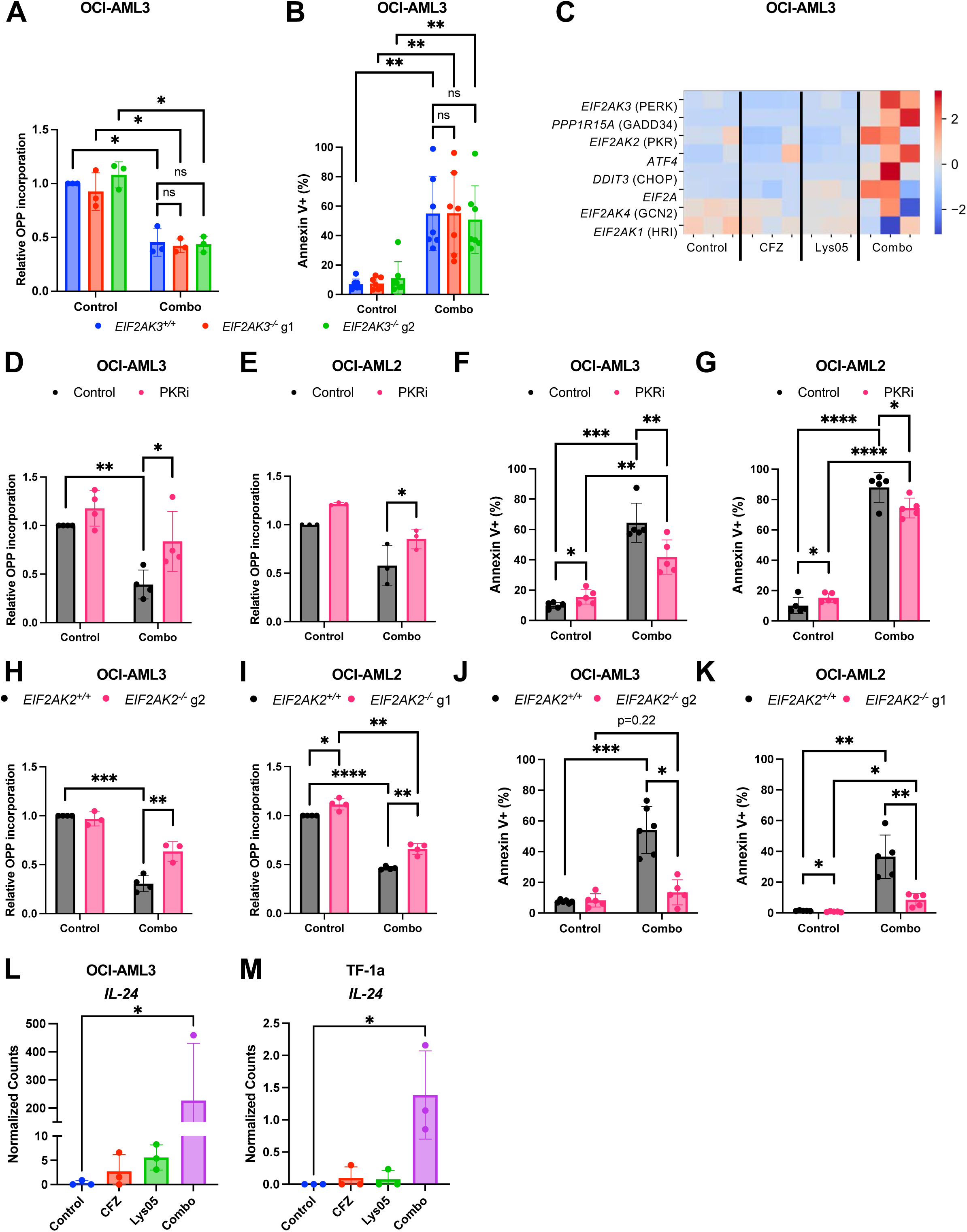
PKR promotes ISR activation in AML cells in response to combined proteasome and autophagy inhibition. (A) Relative OPP incorporation (protein synthesis; n=3) and (B) frequency of apoptotic Annexin V^+^ cells (n=7) in wildtype (*EIF2AK3*^+/+^) and *EIF2AK3*^-/-^ OCI-AML3 cells treated for 24h with vehicle (Control), or the combination of carfilzomib and Lys05 at their respective IC_50_ concentrations (Combo). (B) Heatmap depicting the relative expression (based on RNA-sequencing; n=3) of some ISR genes in OCI-AML3 cells treated for 24h with vehicle (DMSO; Control), 10 nM carfilzomib (CFZ), 7.5 µM Lys05, or the combination of carfilzomib and Lys05 (Combo) (D,E) Relative OPP incorporation (protein synthesis; n=3-4) and (F,G) frequency of apoptotic Annexin V^+^ cells (n=5) in (D,F) OCI-AML3 and (E,G) OCI-AML2 cells treated for 24h with vehicle (control) or the combination of bortezomib and Lys05 (Combo) in the presence or absence of the PKR inhibitor PKR-IN-C16 (PKRi). (H,I) Relative OPP incorporation (protein synthesis; n=3-4) and (J,K) frequency of apoptotic Annexin V^+^ cells (n=5-6) in wildtype (*EIF2AK2*^+/+^) or *EIF2AK2*^-/-^ (H,J) OCI-AML3 and (I,K) OCI-AML2 cells treated for 24h with vehicle (control) or the combination of bortezomib and Lys05 (Combo). (L,M) Relative expression of *IL-24* (from RNA-sequencing data; n=3) in (L) OCI-AML3 and (M) TF-1a cells treated for 24h with DMSO (Control), carfilzomib, Lys05, or the combination of carfilzomib and Lys05 (Combo). Data show individual replicates and mean ± SD in (A, B, D-K). Statistical significance was assessed using a two-tailed paired student’s t-test (A, B, D-K) and using DEseq2 with Kruskal-Wallis test (L,M); ∗p≤0.05, ∗∗p≤0.01, ∗∗∗p≤0.001, ∗∗∗∗p≤0.0001.

In the absence of PERK’s requirement, we wondered if ISR activation was driven by other EIF2α kinases. RNA-sequencing from OCI-AML3 cells treated with carfilzomib, Lys05, combined carfilzomib/Lys05, or vehicle (Fig. S6B), revealed that in addition to PERK, *EIF2AK2* (PKR) was preferentially upregulated in AML cells following carfilzomib/Lys05 treatment (Fig. 6C). To test the contribution of PKR, we treated OCI-AML3 and OCI-AML2 cells with vehicle or combined bortezomib/Lys05 in the presence or absence of the PKR inhibitor PKR-IN-C16^77^. Strikingly, PKR inhibition partially but significantly rescued suppression of protein synthesis and induction of apoptosis in AML cells treated with bortezomib/Lys05 (Fig. 6D-G).

To rule out off-target effects, we engineered *EIF2AK2*^-/-^ OCI-AML3 and OCI-AML2 cells (Fig. S6C,D) and examined effects of bortezomib/Lys05 treatment. Suppression of protein synthesis in response to bortezomib/Lys05 treatment was significantly attenuated in PKR-deficient AML cells (Fig. 6H,I). Furthermore, induction of apoptosis in response to bortezomib/Lys05 treatment was almost completely blunted in the absence of PKR (Fig. 6J,K;). These data suggest that combined proteasome/autophagy inhibition drives effects of ISR activation in a largely PKR-dependent manner.

While PERK is known to be activated by proteostasis disruption, PKR is typically activated by viral infection^73,78^, raising the question of how/why it is a driver of ISR effects in the context of combined proteasome/autophagy inhibition in AML. Prior reports^79,80,81^ linked PKR activation to proteostasis disruption through cytoplasmic accumulation of misfolded IL-24. Combined carfilzomib/Lys05 treatment led to significant upregulation of *IL-24* in OCI-AML3 and TF-1a cells (Fig. 6L,M), suggesting that it may be an underlying driver of PKR activation.

These data suggest that ISR activation proceeds through distinct kinases when proteasome inhibitors are combined with either *HSF1*-deficiency or autophagy inhibition, and that these effects are likely mediated via global and specific changes in proteostasis, respectively.

### Primary Patient AML Stem and Progenitor Cells are Preferentially Sensitive to Proteostasis Disruption

Finally, we tested if combined bortezomib/Lys05 treatment impacted primary patient-derived AML cells (Table S1). Bortezomib/Lys05 treatment suppressed protein synthesis in bulk primary AML cells by ∼50% (Fig. 7A) and increased the frequency of apoptotic cells to ∼75% (Fig. 7B). These effects were also evident within primary AML CD34^+^ stem/progenitor cells, as bortezomib/Lys05 treatment reduced protein synthesis by ∼50% (Fig. 7C) and increased apoptosis to ∼70% (Fig. 7D). In contrast, bortezomib/Lys05 treatment did not significantly alter protein synthesis (Fig. 7E) and only modestly increased apoptosis (Fig. 7F) in normal CD34^+^ hematopoietic stem/progenitor cells isolated from mobilized peripheral blood of healthy adult patients. These data indicate that primary AML cells, including CD34^+^ leukemic stem/progenitor cells, are much more sensitive to combined bortezomib/Lys05 treatment compared to healthy hematopoietic stem/progenitor cells. This suggests that a therapeutic window may exist to preferentially target AML cells, including AML stem/progenitor cells, while largely sparing normal hematopoietic cells.

**Figure 7.**
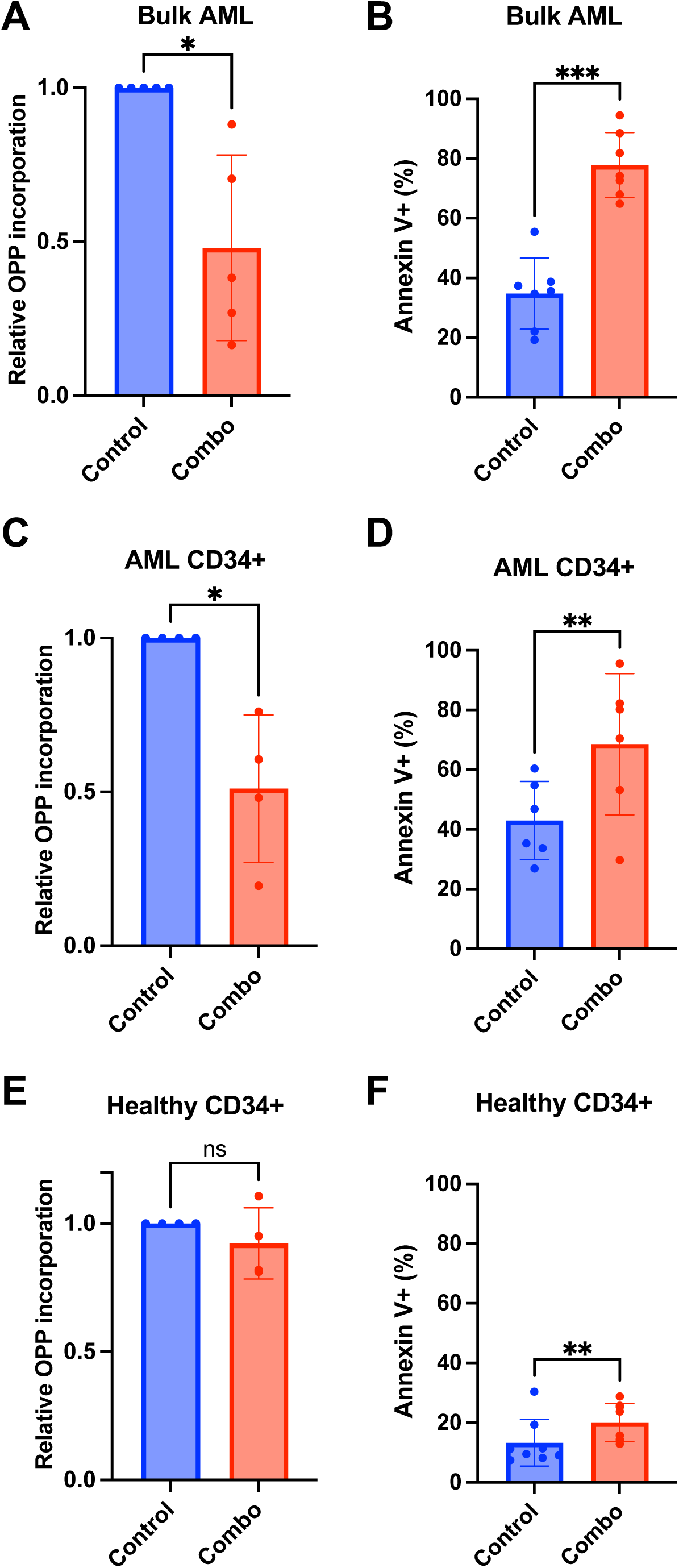
Primary Patient AML Stem and Progenitor Cells are Preferentially Sensitive to Proteostasis Disruption. (A) Relative OPP incorporation (protein synthesis; n=5) and (B) frequency of apoptotic Annexin V^+^ cells (n=7) in primary patient bulk AML cells treated for 24h with vehicle (DMSO; Control) or the combination of 10nM bortezomib and 7.5µM Lys05 (Combo). (B) Relative OPP incorporation (protein synthesis; n=4) and (D) frequency of apoptotic Annexin V^+^ cells (n=6) in primary patient CD34^+^ AML cells treated for 24h with vehicle (DMSO; Control) or the combination of 10nM bortezomib and 7.5µM Lys05 (Combo). (E) Relative OPP incorporation (protein synthesis; n=4) and (F) frequency of apoptotic Annexin V^+^ cells (n=8) in primary healthy CD34^+^ hematopoietic stem and progenitor cells treated for 24h with vehicle (DMSO; Control) or the combination of 10nM bortezomib and 7.5µM Lys05 (Combo). Data show individual replicates and mean ± SD. Statistical significance was assessed using a two-tailed paired student’s t-test; ∗p≤0.05, ∗∗p≤0.01, ∗∗∗p≤0.001.

## DISCUSSION

Here, we uncovered a basis for AML resistance to proteasome inhibition and identified multiple strategies for disrupting proteostasis in AML. In contrast to proteasome inhibitor-sensitive myeloma, bortezomib treatment fails to cause accumulation of misfolded protein in AML cells. Proteostasis in AML is preserved in response to proteasome inhibition at least in part due to activation of HSF1 and increased autophagy. Deletion of *HSF1* or inhibition of autophagy sensitizes AML to proteasome inhibition, leading to accumulation of unfolded protein and ISR activation accompanied by severe declines in protein synthesis, proliferation and survival *in vitro*, and decreased leukemia burden and extended survival *in vivo*.

This study provides key insights into our understanding of proteostasis as a therapeutic target in cancer. Many drugs targeting protein synthesis^82^, folding^16,83^ and degradation^11,23^ have been developed, but their clinical benefits have been modest^16,25^. Our study demonstrates that proteostasis in cancer is regulated by a dynamic and malleable network. Targeting one pathway is typically insufficient to disrupt proteostasis due to activation of adaptive pathways. By disabling these adaptations, proteasome inhibition disrupted proteostasis, leading to severe declines in AML growth, viability and progression. Thus, effective therapeutic strategies will need to consider the proteostasis network as a whole rather than targeting individual pathways.

Proteostasis is highly-conserved, raising hopes that proteostasis-targeting therapies could function across a broad spectrum of cancer types and mutations. AML is genetically heterogeneous^84^ and our studies across four human AML cell lines and seven primary samples represent some of that heterogeneity. Proteostasis disruption impaired each of these AMLs regardless of underlying mutations, suggesting this approach may be effective in a somewhat mutation-agnostic manner. Future studies should focus on understanding how distinct mutations impact proteostasis and proteostasis-targeted therapies. Furthermore, determining whether similar adaptive mechanisms confer resistance to proteasome inhibitors in myeloma specifically, and other cancers generally, could yield significant opportunities to extend use of this class of approved therapeutics to other cancers.

The proteostasis network is configured within HSCs to promote self-renewal^38–41,85–87^. We found that AML cells co-opt proteostasis pathways that are preferentially utilized by HSCs. HSCs are particularly sensitive to proteostasis disruption^38,41^, raising concerns that proteostasis-targeting therapies may impair HSCs. Disrupting proteostasis suppressed protein synthesis and induced apoptosis much more potently in AML stem/progenitor cells than in normal stem/progenitor cells, and treatment regimens were well-tolerated in xenograft recipients. This suggests that therapies aimed at disrupting proteostasis could preferentially kill/impair AML stem/progenitor cells while sparing HSCs and healthy tissues.

Proteasome inhibition combined with *HSF1* deletion led to ISR activation in a PERK-dependent manner. In contrast, combined proteasome/autophagy inhibition led to ISR activation in a PKR-dependent manner. While PERK is canonically tied to proteostasis disruption^76^, PKR is not^73,78^. Recent studies linked PKR activation to proteostasis disruption via accumulation of misfolded IL-24^81^, and we observed increased *IL-24* expression in AML cells in response to carfilzomib/Lys05 treatment. Whether IL-24 contributes to PKR activation in this context should be directly tested in future studies.

Lys05 is a bivalent aminoquinoline that can impair autophagy at lower concentrations than chloroquine or hydroxychloroquine, raising hopes that it can inhibit autophagy in patients^67,88^. It functions by deacidifying lysosomes^88^. While we observed the autophagy inhibitory effects of Lys05, one limitation is that we cannot fully exclude contribution from other lysosomal functions when Lys05 is combined with proteasome inhibition in AML. Future studies could address this by utilizing alternative autophagy inhibitors with distinct mechanisms or through genetic inactivation of autophagy.

In addition to the heat shock response, HSF1 can activate other transcriptional programs^89–91^. Loss of *HSF1* had potent anti-AML effects *in vivo*, consistent with another recent study^45^. Although we cannot fully parse apart the entirety of its impact in AML, we demonstrated that HSF1 has a key role in preserving AML proteostasis and preventing ISR activation.

This study revealed how dynamic configuration of the proteostasis network promotes AML growth and confers resistance to proteasome inhibition. AML cells activate compensatory proteostasis pathways to prevent accumulation of unfolded proteins. Disabling those responses sensitizes AML cells to proteasome inhibition by causing proteostasis disruption and ISR activation. Thus, proteostasis, as a network, is a therapeutic target in AML.

## METHODS

### Cell Culture and Viral Transduction

TF-1a, RPMI 8226, U266, and NCI-H929 cells were cultured in RPMI-1640 (Gibco). MV4-11, OCI-AML2, and OCI-AML3 were cultured in Iscove’s Modified Dulbecco’s Medium (Gibco). HEK293T cells were cultured in Dulbecco’s Modified Eagle Medium (Gibco). Media were supplemented with 10-15% fetal bovine serum (FBS, Gibco) and 1% (v/v) L-glutamine (Corning). NCI-H929 cells were also cultured in 0.05mM 2-mercaptoethanol (Sigma). Cell lines were handled according to American Type Culture Collection protocols.

Transduction of cell lines were performed following a previously described protocol^92^. Briefly, HEK293T cells were grown in 10cm plates and transfected with 10µg of lentiviral plasmid of interest along with 7.5µg psPAX2 (Addgene #12260) and 5µg pMD2.G (Addgene #12259) using polyethylenimine (Polysciences 23966-1). Media was replaced the following day. Supernatant containing viral particles was added to cells with 4µg/ml polybrene (Sigma) and centrifuged at 2000 rpm at 32°C for 90 min.

### Drug Compounds

Bortezomib was purchased from Cell Signaling (#2204) or Bio-Techne (#7282), reconstituted in DMSO and stored at 1mM. Carfilzomib (#S2853), Lys05 (#S8369), and PKR-IN-C16 (#S9668) were purchased from SelleckChem and reconstituted at 10mM in DMSO, 3mg/ml in PBS, and 10mM in DMSO, respectively. All stock solutions were stored in −20°C or −80°C until use.

Due to supply issues, some experiments were performed with either bortezomib or carfilzomib (at empirically determined concentrations for each). In most cases, when supply issues were resolved, independent experiments were performed with both proteasome inhibitors and results were largely similar and reproducible regardless of which proteasome inhibitor was used.

### Proteasome Activity

Proteasome activity was measured using Proteasome-Glo Chymotrypsin-Like Cell-Based Assay (Promega) per manufacturer’s protocol. Bioluminescence was measured using a Tecan Spark plate reader.

### Unfolded Protein

0.5-2×10^6^ cells were treated with the control or the indicated concentrations of drug compounds for 24h. Cells were washed twice with PBS. Tetraphenylethene maleimide (TMI; stock 2mM in DMSO) was diluted in PBS (50µM final concentration) and added to each sample. Samples were incubated with TMI at 37°C for 45 min as previously described^39,41^. Samples were washed twice in PBS and analyzed by flow cytometry (BD Symphony A1).

### RNA-Sequencing

Total RNA was extracted using the RNeasy Plus Micro Kit (Qiagen). Illumina mRNA libraries were prepared using the SMARTseq2^93^ protocol and sequenced in two lanes on the Illumina HiSeq 2500 by the Sequencing core at the La Jolla Institute of Immunology. RNA-sequencing analysis was conducted using the Galaxy platform (usegalaxy.org). Quality control of the raw reads was performed using FastQC^94^, and the reads were trimmed using TrimGalore^95^ and aligned against the human genome assembly GRCh38 using HISAT2^96^. Gene counts were subsequently obtained using featureCounts^97^, and normalization and differentially expressed genes were obtained using DESeq2 (v1.40.2)^98^. P values for differential expression were calculated using the Wald test for differences between the base means of two conditions, then adjusted for multiple test correction using Benjamini Hochberg algorithm. Gene set enrichment analyses (GSEA) were performed on the normalized counts output of DESeq2 using the GSEA software (version 4.3.2) obtained from the Broad Institute (http://www.broadinstitute.org/gsea) and the gene sets from MSigDB version 2023.2. Pathway analyses were performed using Qiagen Ingenuity Pathway Analysis^99^.

### Western Blots

Cell lysates were prepared using RIPA buffer (Thermo Scientific) and protease inhibitor cocktail (Roche). Protein concentrations were quantified using BCA protein assay kit (Thermo Scientific). Lithium dodecyl sulfate loading buffer (Thermo Scientific) was added to proteins and heated at 98°C for 5 min. Between 10-40µg of protein lysate were separated on 4-12% Bis-Tris gels (Life Technologies) and transferred to polyvinylidene fluoride membranes (Bio-Rad). Blots were washed and blocked with 5% bovine serum albumin (Sigma) before being incubated in primary and then secondary antibodies. Blots were developed with the SuperSignal West Pico PLUS, Dura, or Femto chemiluminescent substrate kits (Thermo Scientific) and stripped with 1% SDS, 25mM glycine (pH 2) as needed.

HSF1 (D3L8I; Cell Signaling #12972), phospho^Ser326^-HSF1 (EP1713Y; Abcam #76076), HSP70 (Cell Signaling #4872), GAPDH (14C10; Cell Signaling #2118), phospho^Ser51^-EIF2ɑ (D9G8; Cell Signaling #3398), EIF2ɑ (D7D3; Cell Signaling #5324), ATF4 (D4B8; Cell Signaling #11815), CHOP (L63F7; Cell Signaling #2895), PERK (D11A8; Cell Signaling #5683), PKR (Cell Signaling #3072), LC3B (Cell Signaling #2775), HRP-linked anti-rabbit IgG (Cell Signaling #7074), and HRP-linked anti-mouse IgG (Cell Signaling #7076) antibodies were used for probing.

### GFP-LC3-RFP autophagic flux

Transduction of AML cell lines was performed as described above using pMRX-IP-GFP-LC3-RFP (Addgene #84573)^59^. Cells were treated with the control or the indicated concentrations of carfilzomib or bortezomib and were analyzed by flow cytometry (BD Symphony A1) after 24h.

The mean fluorescence intensity (MFI) of GFP was divided by the MFI of RFP signal to obtain a GFP/RFP ratio.

### CRISPR-Cas9 mediated gene editing

AML cell lines were transduced as described above using lentiCRISPRv2-neo (Addgene #98292)^100^ or lentiCRISPRv2-GFP (Addgene #82416)^100^ cloned with the gRNAs of interest. AML cell lines transduced using lentiCRISPRv2-neo were cultured with 1mg/ml Geneticin (Thermo Scientific) for 7 days and single cells were sorted by fluorescence-activated cell sorting (FACS) to generate individual clones. For the AML cell lines transduced with lentiCRISPRv2-GFP, single GFP-positive cells were sorted by FACS 48-72 hours later to isolate individual clones.

*HSF1*^-/-^ TF-1a cell lines were generated following a previously described transfection protocol^101^. Briefly, gRNA oligos were cloned into pSpCas9(BB)-2A-GFP (Addgene #48138)^101^. TF-1a cells were transfected with the plasmid and Lipofectamine 3000 (Thermo Scientific), and single GFP-positive cells were sorted by FACS 72 h post-transfection to isolate individual clones.

### gRNA sequences

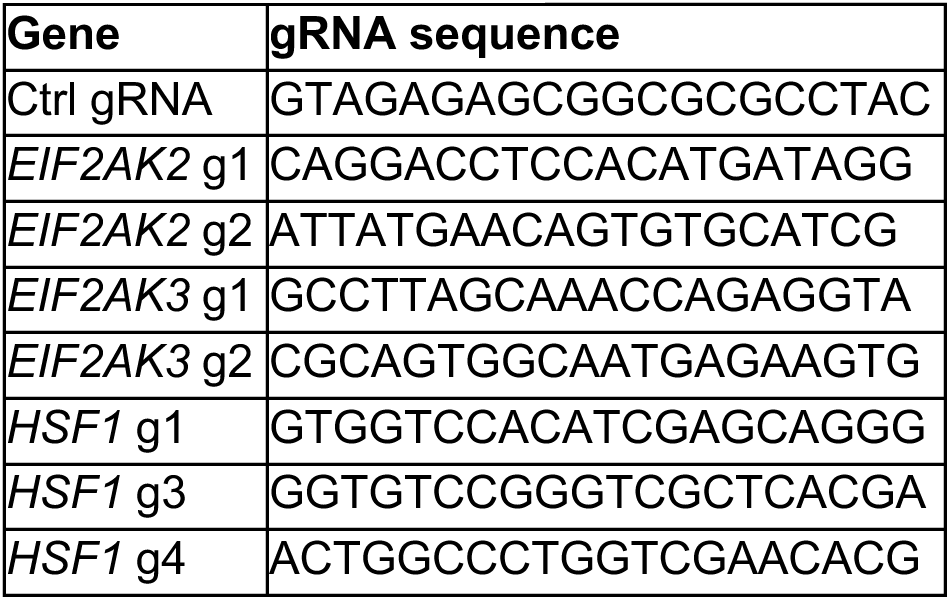

### MTT

Cells were treated with the control (DMSO making up 1% or less of the final concentration of media) or the indicated concentrations of drug compounds for 24h. Then 3-(4,5-dimethyl thiazol- 2-yl)-2,5-diphenyl tetrazolium bromide (MTT) labeling reagent (Roche) was added followed by solubilization buffer 2h later. After overnight incubation, absorbance at 570nm was measured along with 650nm as reference absorbance using a microplate reader (Tecan).

### EdU and OPP

Cells were treated with the control or the indicated concentrations of drug compounds for 24h. Cells were then pulsed with 50-100µM EdU (Thermo Scientific) or 50-100µM OPP (Medchem Source) for 1h, fixed with 1% paraformaldehyde (Thermo Scientific), treated with permeabilization buffer (PBS 1x with 3% (v/v) heat-inactivated bovine serum (Gibco) and 0.1% saponin (Sigma), and processed with Click-iT chemistry (kit from Thermo Scientific) to add AlexaFluor 555 or 647 (Thermo Scientific), and analyzed by flow cytometry (BD Symphony A1) as previously described^40,102^.

### AML Xenografts and Imaging

Transduction of AML cell lines was performed as described above using Lenti-luciferase-P2A-Neo (Addgene #105621)^61^ to generate luciferase-expressing cell lines. Approximately 6 to 10-week-old female NSG mice were transplanted via intravenous tail vein injection with 5×10^5^ luciferase-expressing human AML cell lines. Mice were injected intra-peritoneally (IP) with D-luciferin (150mg/kg bodyweight; PerkinElmer) and luciferase activity was measured using an IVIS200 in vivo imaging system (PerkinElmer) weekly. Bortezomib (0.3-0.5mg/kg retro-orbital intra-venously) and/or Lys05 (30mg/kg IP) were administered weekly over two consecutive days beginning at day 5-7 after transplant until the mice were moribund. All mice were housed in a vivarium at the UC San Diego Moores Cancer Center or at the Sanford Consortium for Regenerative Medicine in specific pathogen-free conditions. All protocols were approved by the UC San Diego Institutional Animal Care and Use Committee.

### Annexin V

Cells were treated with the control or the indicated concentrations of drug compounds for 24h. Cells were then washed with PBS and incubated with FITC- or AlexaFluor 647-tagged Annexin V and propidium iodide (BD Pharmingen™ FITC Annexin V Apoptosis Detection Kit I) for 15-60 min at room temperature prior to being analyzed by flow cytometry (BD Symphony A1).

### RNA interference

4×10^5^ cells were transfected using Lipofectamine RNAiMAX (Thermo Scientific) with 10nM RNAi duplex and 0.2% (v/v) RNAiMAX following the manufacturer’s protocol. 4 siRNAs were pooled for siCtrl. Cells were replated in fresh cell media 20 h post-transfection and cultured for 48h prior to downstream experiments.

### siRNA sequences

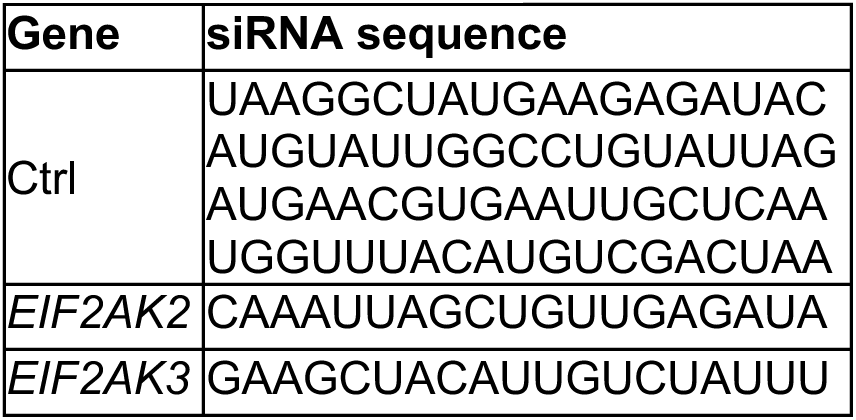

### Primary Patient Samples

Peripheral blood and bone marrow specimens were collected from AML patients at UCSD. Usage of primary samples were approved by the UCSD Institutional Review Board (#180483). Healthy donor specimens were collected by apheresis from healthy allogeneic donors and processed by the UCSD Stem Cell Processing Laboratory. Samples were kept frozen in liquid nitrogen until use. After thawing, samples underwent selection for CD34-positivity using a CD34 MicroBead Kit (Miltenyi Biotech) and cultured in StemSpan SFEM (STEMCELL Technologies) supplemented with 100ng/ml hTPO, 100ng/ml hSCF, 100ng/ml hFLT3L, and 20ng/ml hIL-6 (all cytokines from Peprotech) for 1-2 days prior to use for downstream assays.

### Quantification and Statistical Analysis

Data are represented by mean ± standard deviation (SD) and the statistical test used for each figure panel is described in the figure legends. We performed multiple independent experiments to ensure the reproducibility of our findings as indicated in the figure legends. Mice that died during anesthesia or treatment were excluded from any experiment.

Synergy analyses were performed using Compusyn software (the Chou-Talalay method^68^) and with SynergyFinder+^69^ to calculate combination indexes and synergy scores, respectively.

## AUTHOR CONTRIBUTIONS

K.L., Y.J.K., and R.A.J.S. conceived the project, designed experiments, and wrote the manuscript. K.L. and Y.J.K. performed and analyzed all experiments. C.M.O., A.Z.L., F.J.Z., M.J.S., B.A.C, and S.V provided technical support. J-H.Z, P.W.F., Y.H., E.J.B., L.A.C, and E.D.B. provided key reagents.

## ACKNOWLEDGEMENTS

We thank all patients for providing their blood and/or bone marrow samples in this study. We also thank D.-E. Zhang and C.H.M. Jamieson for reagents and J. Magee for advice. K.L. is supported by NIH/NCI (T32CA121938), NIH/NCATS (KL2TR001444), and an American Society of Hematology Scholar Award. F.J.Z. is supported by NIH/NHLBI (F31HL170531) and NIH/NCI (2T32CA067754). Work in the Signer Laboratory is supported by the NIH/NIDDK (R01DK116951; R01DK124775); NIH/NCI (U01CA267031); the Blood Cancer Discoveries Grant program (8025-20) through The Leukemia & Lymphoma Society, The Mark Foundation for Cancer Research and The Paul G. Allen Frontiers Group; the Cancer Stem Cell Consortium supported by the American Cancer Society and the Lisa Dean Moseley Foundation (CSCC-RSG-23-994830-01-CSCC). K.L. and R.A.J.S are both supported by a Curebound Targeted Grant. K.L., L.A.C and R.A.J.S are supported by the UC San Diego Sanford Stem Cell Institute, Sanford Stem Cell Discovery Center, and the UCSD Moores Cancer Center (NIH/NCI P30CA023100). R.A.J.S and E.J.B. are supported by the NIH/NIA (R01AG088725). S.V. is supported by a CIRM fellowship (EDUC4-12804). L.A.C. is a Scholar of The Leukemia & Lymphoma Society and is supported by NIH/NCI (R37CA252040). Y.H. is supported by the Australian Research Council (FT210100271). The UCSD Moores Cancer Center flow cytometry facility obtained a BD FACSymphony S6 through support from the NIH (S10OD032316). Some figures were generated using Biorender.

## SUPPLEMENTARY INFORMATION

### SUPPLEMENTARY FIGURE LEGENDS

**Supplementary Figure S1 (Related to Figure 1).**
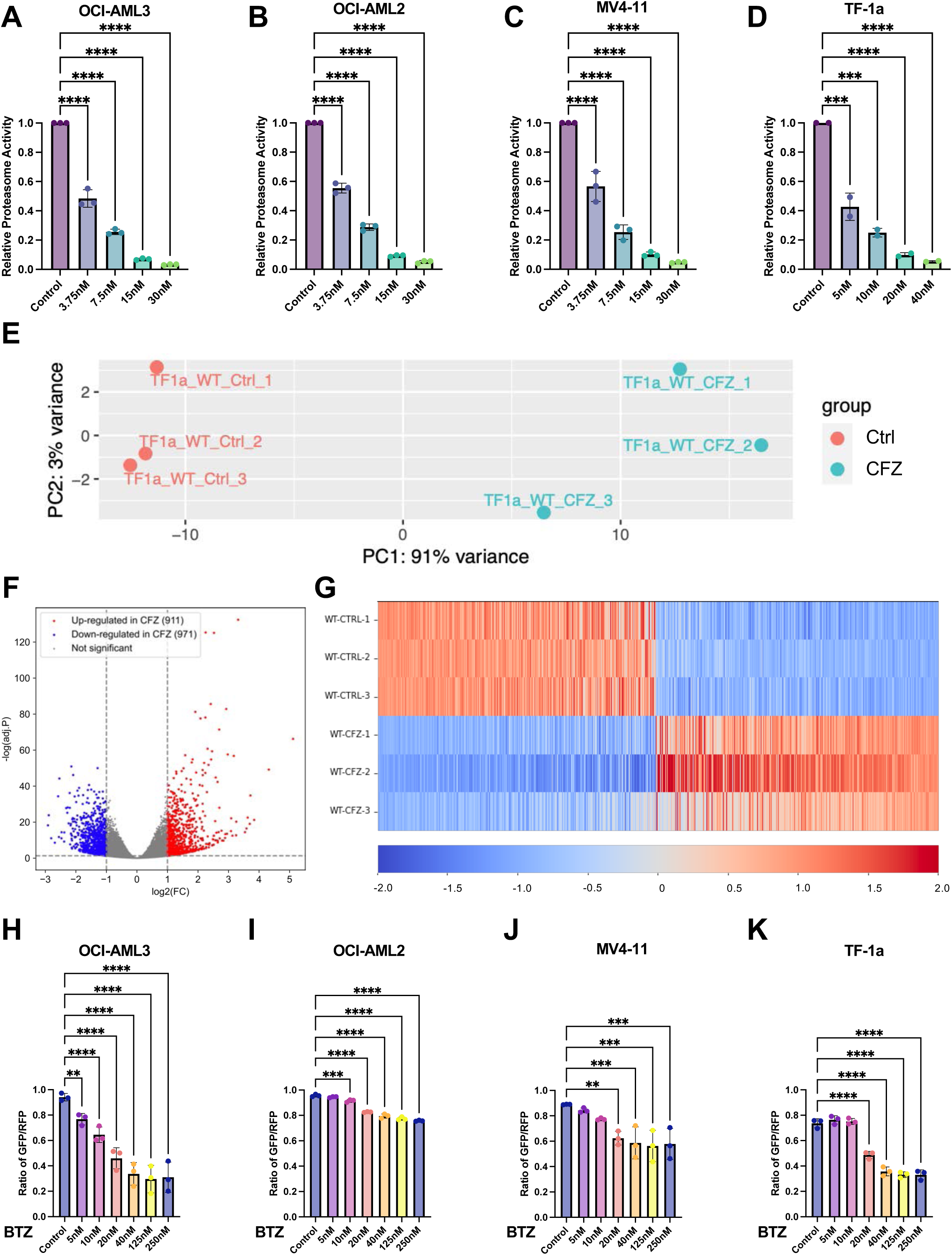
AML Cells Activate Adaptive Proteostasis Pathways in Response to Proteasome Inhibition. (A-D) Relative proteasome activity in (A) OCI-AML3, (B) OCI-AML2, (C) MV4-11, and (D) TF-1a human AML cells lines treated with vehicle (Control) or bortezomib (BTZ) at the indicated concentrations (n=2-3). (E) Principal component analysis of RNA-sequencing data (n=3) demonstrating transcriptomic variation in TF-1a cells treated for 24h with vehicle (Ctrl) or 125nM carfilzomib (CFZ). (F) Volcano plot depicting significantly upregulated (red) and downregulated (blue) expression of genes in TF-1a cells treated for 24h with 125nM carfilzomib (CFZ) as compared to vehicle treated controls. Dashed lines indicate the statistical cutoff of |log2(FC)|>1 and adjusted p-value < 0.05. (G) Heatmap depicting the most differentially expressed genes in TF-1a cells treated for 24h with vehicle (Ctrl) or 125nM carfilzomib (CFZ). (H-K) Ratio of GFP:RFP fluorescence (inversely correlated to autophagic flux) measured by flow cytometry in (H) OCI-AML3, (I) OCI-AML2, (J) MV4-11, and (K) TF-1a human AML cell lines transduced with the RFP-GFP-LC3 reporter and treated for 24h with bortezomib (BTZ) at the indicated concentrations (n=3). Data show individual replicates and mean ± SD in (A-D, H-K). Data were assessed using ANOVA followed by Dunnett’s test relative to control (A-D, H-K). ∗p≤0.05, ∗∗p≤0.01, ∗∗∗p≤0.001, ∗∗∗∗p≤0.0001.

**Supplementary Figure S2 (Related to Figure 2).**
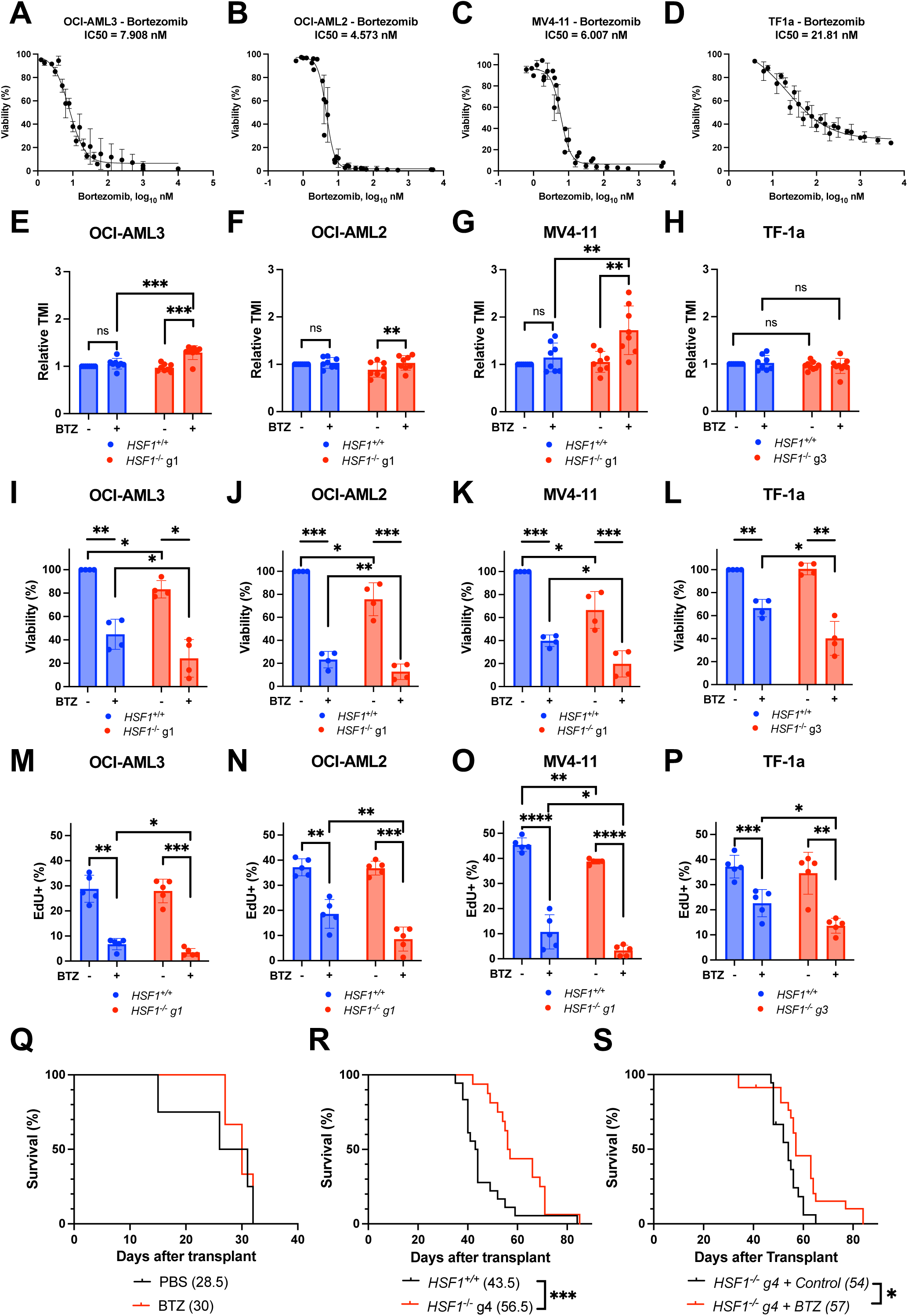
*HSF1* deletion sensitizes human AML cells to proteasome inhibition *in vitro* and *in vivo*. (A-D) Frequency of viable cells (measured by MTT) in (A) OCI-AML3 (n=8), (B) OCI-AML2 (n=6), (C) MV4-11 (n=6), and (D) TF-1a (n=8) human AML cells lines cultured for 24h in the presence of increasing concentrations of bortezomib. IC_50_ concentrations calculated based on these curves are shown. (E-H) Relative TMI fluorescence (unfolded protein content) detected by flow cytometry in wildtype (*HSF1*^+/+^) and *HSF1*^-/-^ (E) OCI-AML3, (F) OCI-AML2, (G) MV4-11, and (H) TF-1a human AML cell lines treated for 24h with vehicle (PBS; -) or their respective IC_50_ concentrations of bortezomib (BTZ; +; n=8). (I-L) Frequency of viable cells (measured by MTT) in wildtype (*HSF1*^+/+^) and *HSF1*^-/-^ (I) OCI-AML3, (J) OCI-AML2, (K) MV4-11, and (L) TF-1a human AML cell lines treated for 24h with vehicle (PBS; -) or their respective IC_50_ concentrations of bortezomib (BTZ; +; n=4). (M-P) Frequency of dividing EdU^+^ (following 1h pulse) wildtype (*HSF1*^+/+^) and *HSF1*^-/-^ (M) OCI- AML3, (N) OCI-AML2, (O) MV4-11, and (P) TF-1a cells treated with vehicle (PBS; -) or their respective IC_50_ concentrations of bortezomib (BTZ; +; n=5). (Q) Overall survival of NSG mice xenografted with wildtype TF-1a cells and treated with PBS (n=4) or 0.5 mg/kg bortezomib (BTZ; n=3). Treatment was initiated 5 days after transplant and drug was administered weekly for 2 consecutive days followed by 5 days with no treatment. (R) Overall survival of NSG mice xenografted with wildtype (n=18) or *HSF1*^-/-^ (n=16) TF-1a cells (transduced with luciferase). (S) Overall survival of NSG mice xenografted with *HSF1*^-/-^ TF-1a cells (transduced with luciferase) and treated with PBS (n=18) or 0.5 mg/kg bortezomib (BTZ; n=23). Treatment was initiated 5 days after transplant and drug was administered weekly for 2 consecutive days followed by 5 days with no treatment. Data show individual replicates and mean ± SD in (E-P). Statistical significance was assessed using a two-tailed paired student’s t-test (E-P) or log-rank test (Q-S); ∗p≤0.05, ∗∗p≤0.01, ∗∗∗p≤0.001, ∗∗∗∗p≤0.0001.

**Supplementary Figure S3 (Related to Figure 3).**
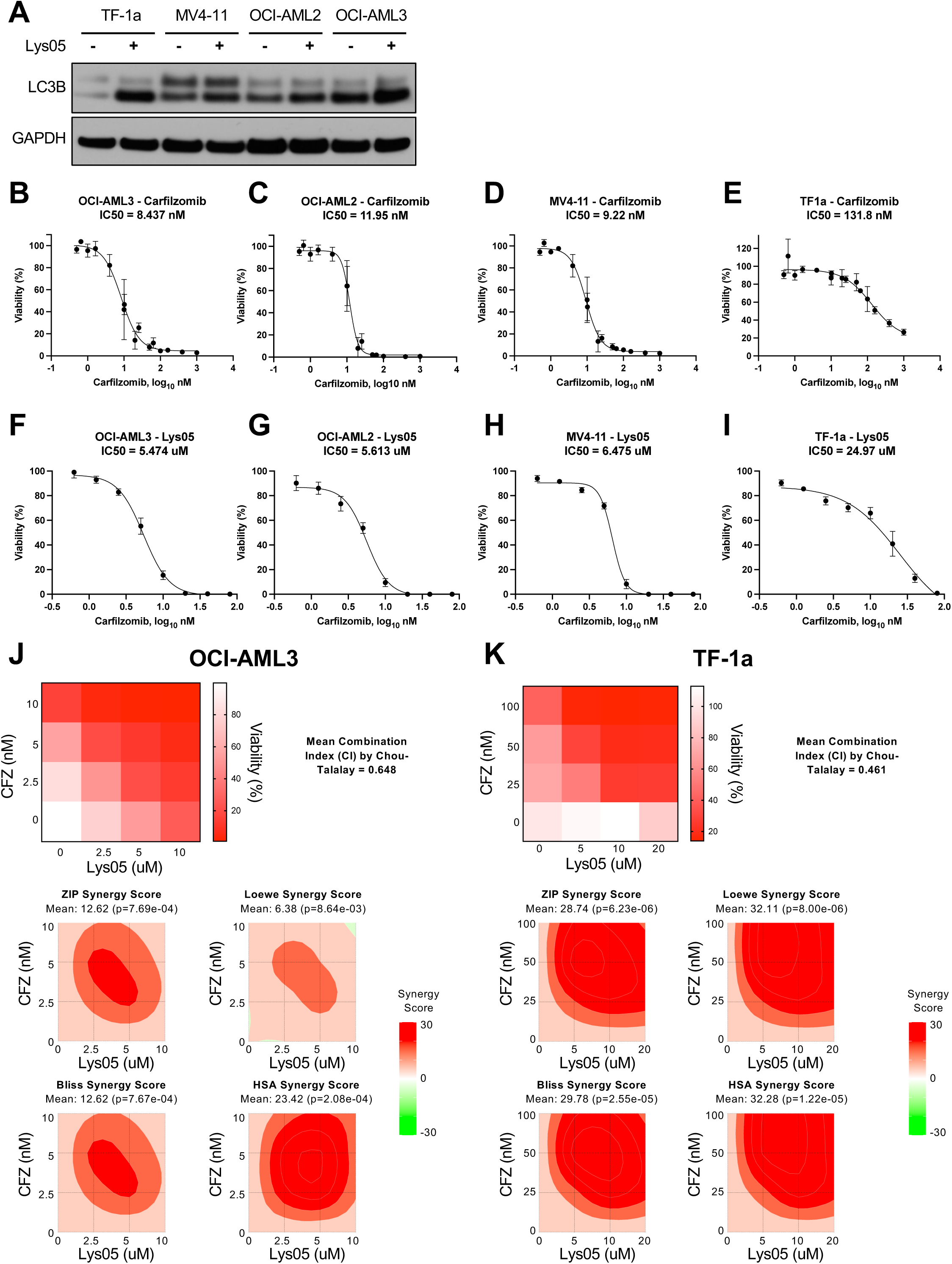
Combined proteasome and autophagy inhibition synergistically disrupts AML proteostasis, proliferation and survival *in vitro* and impairs AML progression *in vivo*. (A) Western blot for LC3B expression in TF-1a, MV4-11, OCI-AML2, and OCI-AML3 human AML cell lines treated for 4h with vehicle (PBS; -) or their respective IC_50_ concentrations of Lys05 (+). One of 2 representative blots shown. (B-E) Frequency of viable cells (measured by MTT) in (B) OCI-AML3, (C) OCI-AML2, (D) MV4-11, and (E) TF-1a human AML cells lines cultured for 24h in the presence of increasing concentrations of carfilzomib (n=6). IC_50_ concentrations calculated based on these curves are shown. (F-I) Frequency of viable cells (measured by MTT) in (F) OCI-AML3, (G) OCI-AML2, (H) MV4-11, and (I) TF-1a human AML cells lines cultured for 24h in the presence of increasing concentrations of Lys05 (n=3). IC_50_ concentrations calculated based on these curves are shown. (J-K) Frequency of viable (measured by MTT) (J) OCI-AML3 and (K) TF-1a cells treated with varying concentrations of carfilzomib and Lys05. The mean combination index using the Chou-Talalay method via Compusyn^68^ as well as synergy plots produced using SynergyFinder+^69^ for zero interaction potency (ZIP), Loewe, Bliss, and highest single agent (HSA) synergy models with synergy scores are shown. Data show individual replicates and mean ± SD in (B-I).

**Supplementary Figure S4 (Related to Figure 4).**
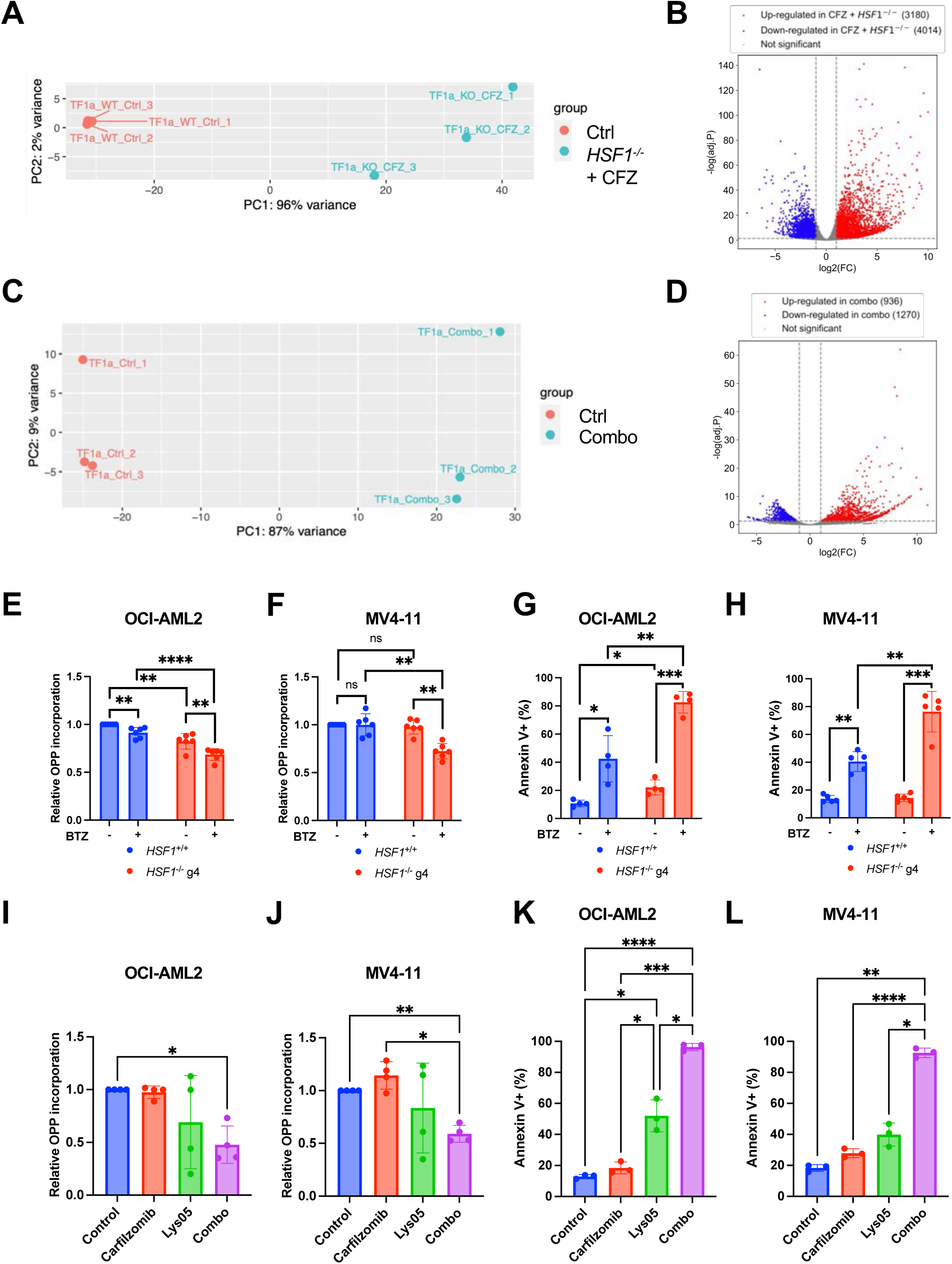
Proteostasis disruption activates a terminal ISR in human AML cells. (A) Principal component analysis of RNA-sequencing data (n=3) demonstrating transcriptomic variation in vehicle-treated wildtype TF-1a cells (WT_Ctrl) and *HSF1*^-/-^ TF-1a cells treated for 24h with 125nM carfilzomib (KO_CFZ). (B) Volcano plot depicting significantly upregulated (red) and downregulated (blue) expression of genes in *HSF1*^-/-^ TF-1a cells treated for 24h with 125nM carfilzomib as compared to vehicle-treated wildtype TF-1a cells. Dashed lines indicate the statistical cutoff of |log2(FC)|>1 and adjusted p-value < 0.05. (C) Principal component analysis of RNA-sequencing data (n=3) demonstrating transcriptomic variation in TF-1a cells treated for 24h with vehicle (DMSO; Ctrl) or the combination of 125nM carfilzomib and 20µM Lys05 (Combo). (D) Volcano plot depicting significantly upregulated (red) and downregulated (blue) expression of genes in TF-1a cells treated for 24h with the combination of 125nM carfilzomib and 20µM Lys05 (Combo) as compared to vehicle (DMSO) treated controls. Dashed lines indicate the statistical cutoff of |log2(FC)|>1 and adjusted p-value < 0.05. (E-F) Relative O-propargyl-puromycin (OPP) incorporation (protein synthesis) in wildtype (*HSF1*^+/+^) and *HSF1*^-/-^ (E) OCI-AML2 (n=6) and (F) MV4-11 (n=6) cells treated for 24h with vehicle (PBS; -) or their respective IC_50_ concentrations of bortezomib (BTZ; +). (G-H) Frequency of apoptotic Annexin V^+^ wildtype (*HSF1*^+/+^) and *HSF1*^-/-^ (G) OCI-AML2 (n=4) and (H) MV4-11 (n=5) cells treated for 24h with vehicle (PBS; -) or their respective IC_50_ concentrations of bortezomib (BTZ; +). (I-J) Relative OPP incorporation (protein synthesis) in (I) OCI-AML2 and (J) MV4-11 cells treated for 24h with DMSO (Control), carfilzomib, Lys05, or the combination of carfilzomib and Lys05 (Combo). Carfilzomib and Lys05 were used at their respective IC_50_ concentrations for each cell line (n=4). (K-L) Frequency of apoptotic Annexin V^+^ (K) OCI-AML2 and (L) MV4-11 cells treated for 24h with DMSO (Control), carfilzomib, Lys05, or the combination of carfilzomib and Lys05 (Combo). Carfilzomib and Lys05 were used at their respective IC_50_ concentrations for each cell line (n=3). Data show individual replicates and mean ± SD in (E-L). Statistical significance was assessed using a two-tailed paired student’s t-test (E-H) or ANOVA followed by Tukey’s test (I-L); ∗p≤0.05, ∗∗p≤0.01, ∗∗∗p≤0.001, ∗∗∗∗p≤0.0001.

**Supplementary Figure S5 (Related to Figure 5).**
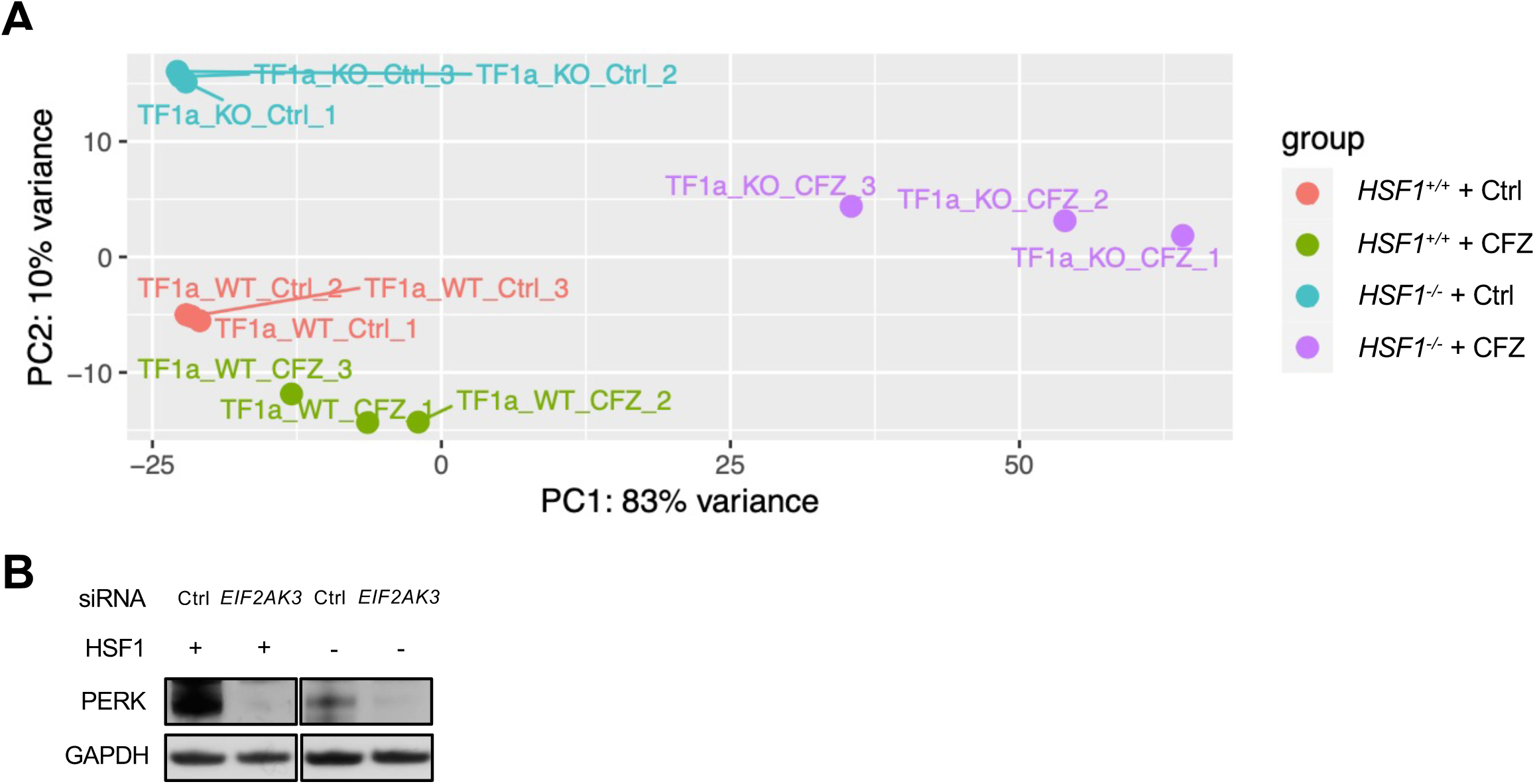
PERK promotes ISR activation in *HSF1*-deficient AML cells in response to proteasome inhibition. (A) Principal component analysis of RNA-sequencing data (n=3) demonstrating transcriptomic variation in wildtype and *HSF1*^-/-^ TF-1a cells treated for 24h with vehicle or 125nM carfilzomib (CFZ). (B) Western blot showing the expression of PERK in wildtype and *HSF1*^-/-^ OCI-AML3 cells 72h after transfection with a control (Ctrl) or *EIF2AK3* targeted siRNA.

**Supplementary Figure S6 (Related to Figure 6).**
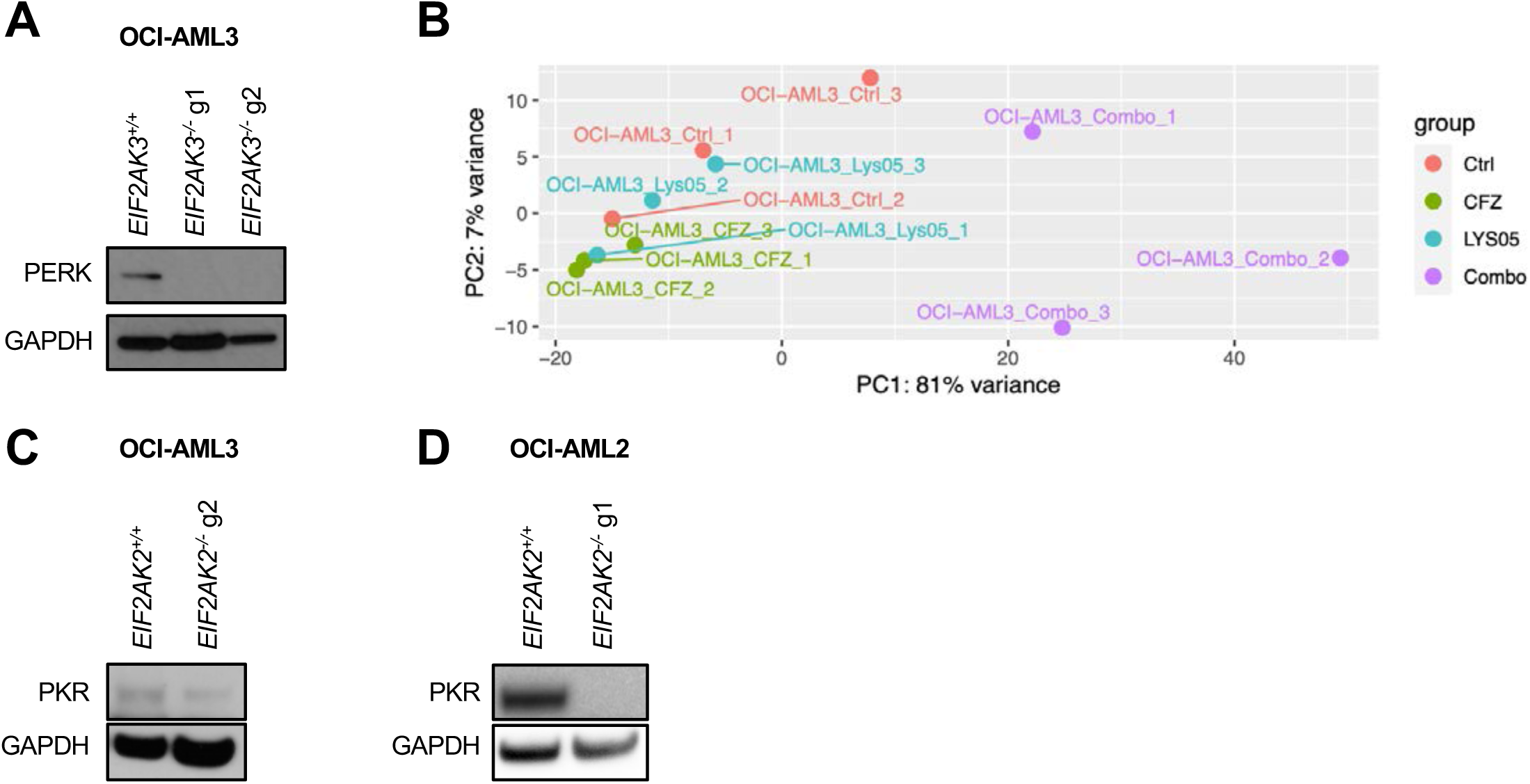
PKR promotes ISR activation in AML cells in response to combined proteasome and autophagy inhibition. (A) Western blot showing expression of PERK from wildtype and *EIF2AK3*^-/-^ OCI-AML3 cells. (B) Principal component analysis of RNA-sequencing data (n=3) demonstrating transcriptomic variation in OCI-AML3 cells treated for 24h with vehicle (DMSO; Ctrl), 10nM carfilzomib (CFZ), 7.5µM Lys05, or the combination of carfilzomib and Lys05 (Combo). (C,D) Western blot showing expression of PKR in wildtype and *EIF2AK2*^-/-^ (C) OCI-AML3 and (D) OCI-AML2 cells.

**Supplementary Table S1. (Related to Figure 7).**
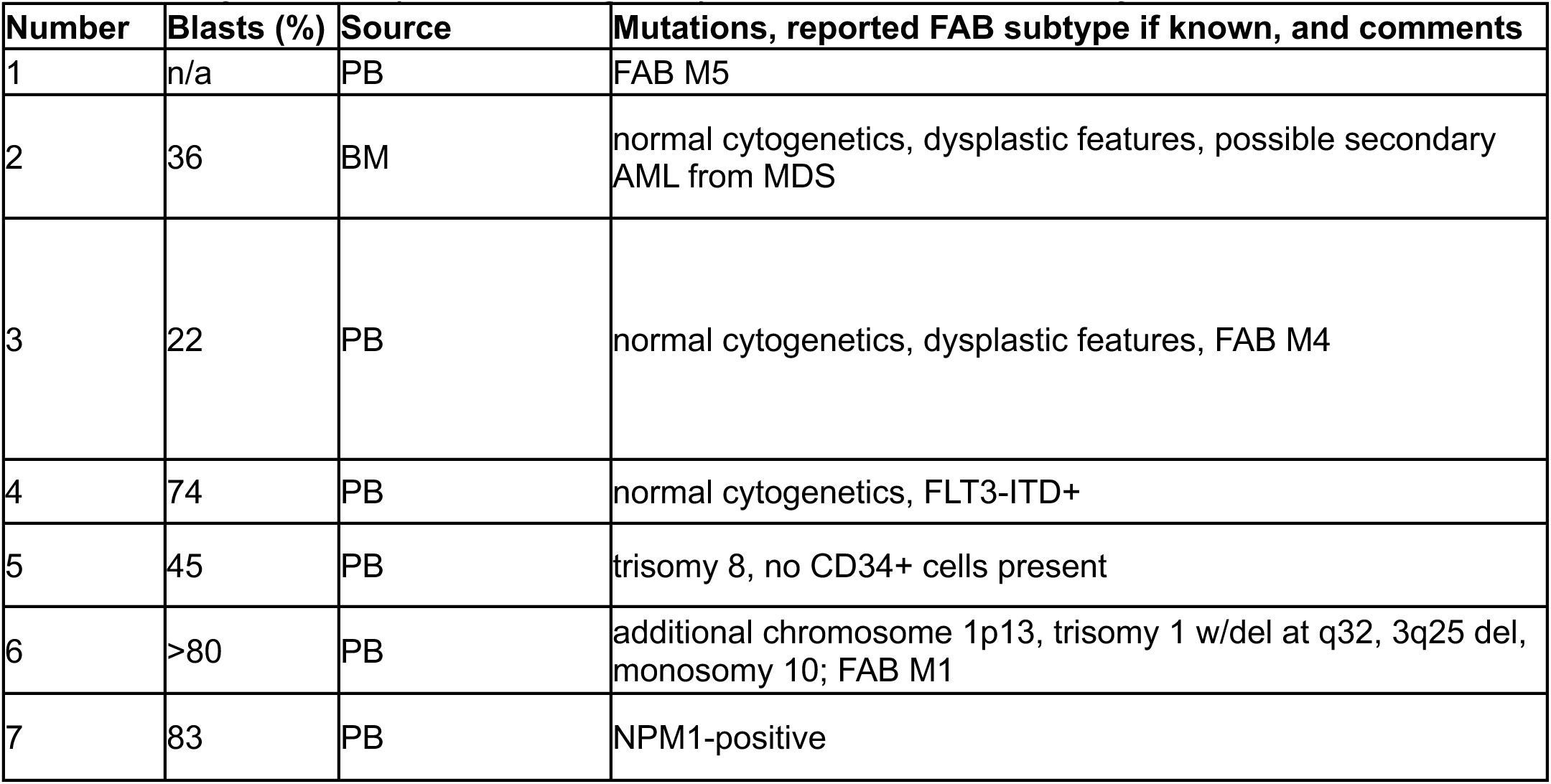
Characteristics of primary AML samples.

| **Number** | **Blasts (%)** | **Source** | **Mutations, reported FAB subtype if known, and comments** |
| --- | --- | --- | --- |
| 1 | n/a | PB | FAB M5 |
| 2 | 36 | BM | normal cytogenetics, dysplastic features, possible secondary AML from MDS |
| 3 | 22 | PB | normal cytogenetics, dysplastic features, FAB M4 |
| 4 | 74 | PB | normal cytogenetics, FLT3-ITD+ |
| 5 | 45 | PB | trisomy 8, no CD34+ cells present |
| 6 | >80 | PB | additional chromosome 1p13, trisomy 1 w/del at q32, 3q25 del, monosomy 10; FAB M1 |
| 7 | 83 | PB | NPM1-positive |

